# Differences in Cellular mechanics and ECM dynamics shape differential development of wing and haltere in *Drosophila*

**DOI:** 10.1101/2024.04.26.591286

**Authors:** C Dilsha, Salima Shiju, Neel Ajay Shah, Mandar M. Inamdar, L S Shashidhara

**Affiliations:** Indian Institute of Science Education and Research, Pune; Indian Institute of Technology Bombay, Mumbai 400076; National Centre for Biological Sciences, Bangalore; Ashoka University, New Delhi

**Keywords:** Haltere, wing, ECM, morphogenesis, Ubx, Atrophin, Pten, Expanded, 3-D tissue shape, cellular mechanical properties, organ shape

## Abstract

Diverse organ shapes and sizes arise from the complex interplay between cellular properties, mechanical forces, and gene regulation. *Drosophila* wing-a flat structure and the globular haltere are two homologous flight appendages emerging from a similar group of progenitor cells. The activity of a single Hox transcription factor, Ultrabithorax (Ubx), governs the development of these two distinct organs-wing and haltere with different cell and organ morphologies. Our work reported here on differential development of wing and haltere suggest that the localisation and abundance of actomyosin complexes, apical cell contractility, properties of extracellular matrix, and cell size and shape, which is a result of various cell intrinsic and extrinsic forces, plausibly influence the flat vs. globular geometry of these two organs. We followed the three-dimensional architecture of developing wing and halteres during early pupal morphogenesis, indicating the role of the above-mentioned factors in force generation and in driving differential morphogenesis, leading to different organ shapes. Loss of Ubx function led to wing cell-like cellular features in haltere discs, and corresponding changes at the level of adult organs. We also observed that RNAi-mediated downregulation of *Atrophin* or *Pten*, in the background of downregulated *Expanded* (or elevated Yki), gave rise to varying degrees of haltere to wing homeotic transformations at the cellular as well as adult organ levels. Finally, we provide simulated scenarios based on computational modelling that propose key ingredients required for producing wing- or haltere-like morphologies in *Drosophila*.

## INTRODUCTION

Specific morphology and dimension of various organs are vital for their functionality and, in turn, for the survival of the organism. The properties of individual cells and their arrangement within developing tissues contribute to their overall architecture and mechanical properties. Epithelial tissues undergo extreme topological changes like folding, bending, and flattening during organogenesis by generating local and global force asymmetries (Nelson & Gleghorn, 2012; Tozluoǧlu & Mao, 2020; Zartman & Shvartsman, 2010). These dynamic changes are crucial for transforming a flat two-dimensional (2D) epithelium into a three-dimensional (3D) structure and contribute to the diversity of organ shapes and functions observed in living organisms. Thus, epithelial cells exhibit remarkable plasticity and are capable of adopting various shapes, arrangements, and structural features during development. This plasticity is facilitated by the dynamic interplay between intracellular signalling pathways, spatiotemporal patterning of cytoskeletal regulators, cell-cell adhesion and interactions between cell and extracellular matrix (ECM) (Díaz-de-la-Loza & Stramer, 2024; Kozyrina et al., 2020; Luciano et al., 2022; Miao & Blankenship, 2020).

The differential development of *Drosophila* wing and haltere offers a unique system to investigate the processes and factors that shape cellular and organ-level morphology and growth. Despite originating from remarkably similar progenitor cells, these two organs ultimately differentiate into distinct functional structures. The adult wing is a flat bi-layered structure composed of squamous cells, whereas the haltere, also bi-layered, is smaller and globular, comprising cuboidal cells (Fig 1A) (Roch & Akam, 2000). Unlike in the wing, the two layers of the haltere—dorsal and ventral—are not closely apposed together and lack veins and interveins, contributing to its morphology distinct from that of wings. The hox gene, *Ultrabithorax* (*Ubx*), which is only expressed in halteres, is capable of specifying all aspects of haltere development from the default wing fate, as the loss of function of *Ubx* in halteres leads to a complete homeotic transformation resulting in a pair of wings instead of halteres (Lewis, 1978). Thus, differential development of wing and haltere is a unique genetic assay system to study mechanisms of by which two different epithelial morphologies are attained.

**Figure 1:**
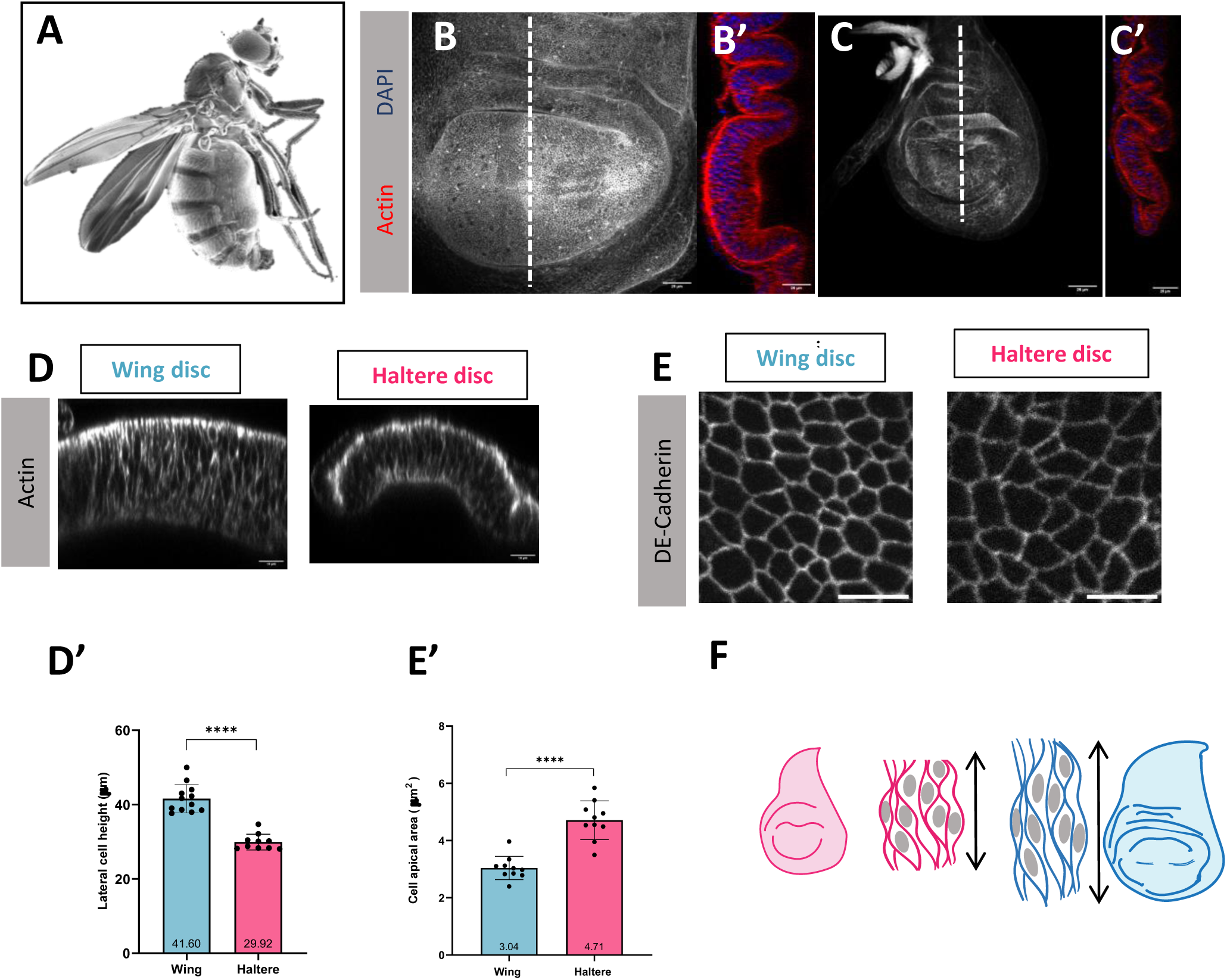
Differential cell apical area and lateral cell height in the columnar cells of larval third instar wing and haltere disc cells. A. Scanning Electron Micrograph of an adult *Drosophila*. Note the adult flat wing and “knob-like” haltere morphology. B. Maximum intensity projection of third instar wing disc stained for actin (grey). B’. Lateral cross-section along the dashed white line shows the organisation of DP cells in the pouch, F-Actin (red) and DAPI (blue). Scale bar, 25 μm. C. Maximum intensity projection of third instar haltere disc stained with actin (grey). C’. Lateral cross-section of haltere disc along the white dashed line shows the organisation of DP cells in the pouch, F-actin (red) and DAPI (blue). Scale bar, 25 μm. D. Cross section of wing and haltere pouch DP cells stained for F-actin (grey) showing the pseudostratified columnar cells. Haltere cells are laterally less elongated compared to the wing disc cells. Scale bar 10 µm. D’. Quantification of the lateral cell height of wing and haltere discs at L3. The mean, s.d and individual data points are presented. **** p<0.0001, Student’s t-test n= 12 and 10 discs for wing and haltere, respectively. E. Apical projection of wing and haltere disc cells stained for *DE-Cadherin*. Wing disc cells are relatively more apically constricted compared to the haltere discs. Scale bar, 5 μm. E’ Quantification of cell apical area. The mean, standard deviation and individual data points are plotted. **** p<0.0001, Student t-test, n=10 discs each for each wing and haltere. F. Illustration showing the differences between the pseudostratified columnar cells of wing and haltere. The cells of the haltere discs, although columnar, exhibit a reduced lateral height and a relaxed apical area compared to the corresponding wing cells at L3.

Wing and haltere discs are composed of columnar cells at the third instar larval stages (L3). The discs undergo dramatic morphogenesis at the pupal stages, transforming the ‘sac-like’ imaginal discs into their respective adult morphologies. Recent studies have shown that wing discs undergo columnar to cuboidal cell shape transition and, apical and basal ECM remodelling, aiding in tight apposition of dorsal and ventral layers at the early hours of puparium formation (7-9h APF) (De las Heras et al., 2018; Diaz-de-la-Loza et al., 2018). Cells adhere to the ECM through transmembrane proteins such as integrins which are linked to the cytoskeleton and transmit mechanical signals bidirectionally (Katsumi et al., 2004; Schwartz, 2010; Z. Sun et al., 2016). During early puparium formation, because of *Ubx* function, the columnar shape of the haltere cells is retained and Matrix metallopeptidases (MMP1 and MMP2) and other proteases, such as Stubble (Sb) and Notopleural (Np), necessary for the matrix remodelling are repressed thereby preventing ECM degradation (De Las Heras et al., 2018; Diaz-de-la-Loza et al., 2020). This retention of cell shape and ECM is thought to be one of the major factors preventing dorso-ventral apposition in halteres and, thereby, maintaining the globular geometry. However, degrading the basal ECM (bECM) in halteres by expressing MMP1 (at early L3) resulted only a mild flattening (De las Heras et al., 2018). Interestingly, inducing ECM degradation at late L3 resulted in observable cell shape changes, but the organs retained their globular morphology. Moreover, it was observed that the downregulation of Ubx at late L3 stages leads to globular halteres instead of flat ‘winglets’ (De las Heras et al., 2018). These findings imply that critical changes may accumulate between wing and haltere discs during or prior to the third instar larval stage, which are essential for shaping their respective morphologies, emphasizing the need to identify additional factors that might play a role during the L3 stage.

Until the third instar larval stages, apart from the difference in cell number and, consequently, tissue size, the wing and haltere disc cells were generally regarded as indistinguishable in terms of their individual epithelial cell size, shape, and polarity (Makhijani et al., 2007; Roch & Akam, 2000; Singh et al., 2015). However, previous studies have reported that *Ubx^-^* clones in the halteres tend to sort out from their surrounding wildtype haltere cells, indicating their differential adhesion properties (Morata and Garcia-Bellido, 1976; Shashidhara et al., 1999). The role of Ubx in modulating various growth and wing patterning pathways has been extensively studied (Agrawal et al., 2011; Crickmore & Mann, 2006; Khan et al., 2020; Makhijani et al., 2007; Mohit et al., 2006; Pallavi et al., 2006; Pavlopoulos & Akam, 2011; Prasad et al., 2003; Shashidhara et al., 1999; Weatherbee et al., 1998). Nevertheless, to form such distinct structures, the various signalling networks and cues from the external environment must converge at the level of altering the individual cell characteristics such as cell shape, size and mechanical properties-dictating the overall tissue geometry. However, our understanding of the mechanisms responsible for shaping organs and how Ubx specifies the globular shape of the haltere remains limited.

With the recent advances in microscopy techniques, quantitative imaging, and analysis, we re-investigated the properties of wing and haltere epithelial cells at L3. Here we report, hitherto unrecognised mechanical properties of the haltere epithelial cells at third instar larval stages, which plausibly are the reasons for adult halteres to attain a morphology different from that of adult wings.

## RESULTS

*Drosophila* wing and haltere imaginal discs have very similar overall morphology until the L3 stage, and the discs become progressively different from each other at subsequent pupal stages (Roch & Akam, 2000). Third instar wing and haltere discs can be physically subdivided into areas of pouch, hinge and notum. They are separated by their characteristic folds and three-dimensional architecture (Fig 1B, B’, C, C’). The cells of the respective pouch region give rise to the adult wing blade and haltere capitellum; hence, our study majorly focuses on the characteristics of the pouch cells in these discs at various stages of their development. Both the wing and haltere discs consist of a single layer of pseudostratified columnar epithelium, also referred to as disc proper (DP), and a squamous epithelium located apically to the DP, which forms the peripodial membrane (Fig 1B’, C’, D). Halteres and their corresponding imaginal discs are smaller than wings, containing fewer number of cells (P. F. Martin, 1982).

### Haltere disc cells are shorter and have larger apical surface area compared to wing cells at L3 stage

We first examined cell size, shape and tissue architecture in wing and haltere discs in L3 stage. To visualise pseudostratified columnar DP cells, we stained the L3 wing and haltere discs with Rhodamine Phalloidin. Although concentrated at the apical belt of the cells, the actin staining was present along the entire length of the columnar cells. This was further confirmed by the co-visualisation of F-actin and a basal membrane marker Viking GFP (Collagen IV α2-subunit, component of basal ECM) (S1A).

Interestingly, we observed differences in lateral cell heights between the cells of wing and haltere discs. We found that the wing disc cells were laterally more elongated and had an average cell height of approximately 40 microns, whereas the haltere disc cells were notably shorter, averaging close to 30 microns (Fig 1D, D’).

We further compared the cell apical surface areas between the wing and haltere discs using *DE-cadherin* as a marker to visualise the apical membrane. We performed cell segmentation on acquired confocal images using Tissue Analyzer (Aigouy et al., 2016). We observed larger apical surface area in haltere cells compared to wing cells (Fig 1E, E’). We also tried to approximate the volume of the pseudostratified columnar cells between wing and haltere cells. Since the precise estimation of the volume of pseudostratified columnar cells is difficult, we approximated the volume by multiplying a known area with the lateral cell height and dividing it by the number of nuclei. Our preliminary data suggest that the volume of the wing and haltere cells are comparable (S1B). Taken together, our results suggest that the columnar cells of wing disc DP are laterally more elongated and apically more constricted compared to halteres discs at the L3 stage (Fig 1F).

### Haltere disc cells shows lower levels of Actin and Myosin II compared to wing disc cells

Actin and Myosin, key components of the cytoskeleton, play pivotal roles in controlling cell size and shape by orchestrating dynamic changes in the cellular architecture. Apical constrictions are predominantly mediated by actin filaments and Myosin-II motors (Heer & Martin, 2017; Kondo & Hayashi, 2015). Since we observed differences in the lateral cell height and apical surface area between the L3 wing and haltere discs, we explored whether these differences stem from variations in the levels of actin and Myosin II accumulation at the apical cortex. We stained wing and haltere imaginal discs with Rhodamine Phalloidin to visualize F-actin. Although actin is distributed throughout the length of the DP cells, it is noticeably concentrated in the apical cortex of both the discs. We quantified the mean fluorescence intensity between the two discs and observed lower levels of actin intensity at the apical cortex of haltere disc cells compared to wing disc cells at L3 (Fig 1D, 2A).

Similarly, we measured the Myosin II levels using the light regulatory chain *Spaghetti squash*/(sqh) using a GFP-tagged transgene (Sqh GFP) line (referred to hereafter as Myosin or Myosin II). The Myosin was densely localised in the peripodial cells and apical cortex of DP cells. However, the DP cells of haltere discs showed lower levels of Myosin in the apical cortex than the wing disc cells at L3 (Fig 2B, C).

**Figure 2.**
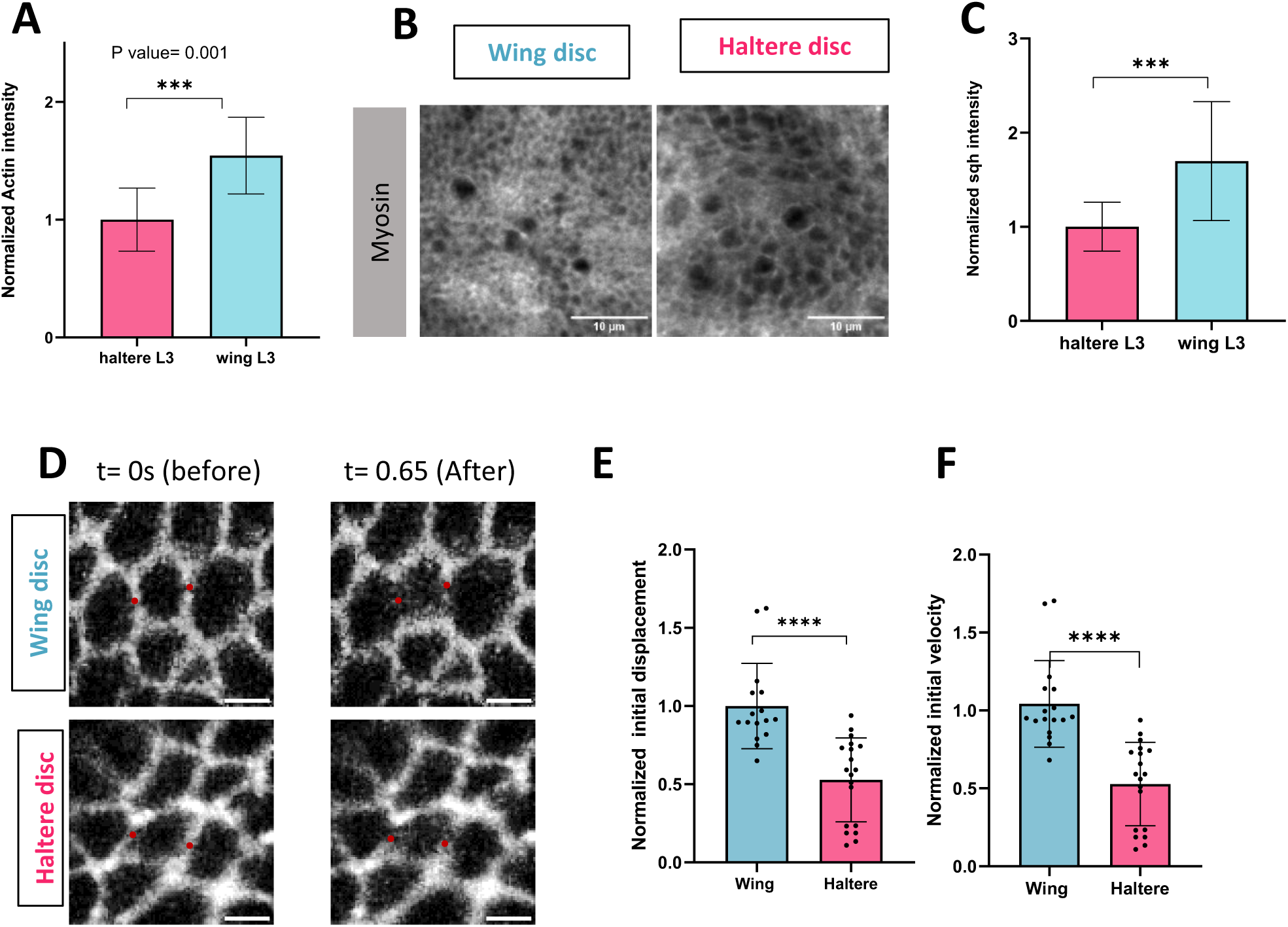
Wing and haltere cells shows differential actomyosin accumulation and junctional tension at cell apical cortex. A. Quantification of F-actin fluorescence intensity at the apical cortex of wing and haltere DP cells at L3. Normalised actin intensity is plotted. The mean and s.d are presented, *** p<0.001, Student’s t-test, n=11 discs for both wing and haltere. B. Myosin localisation at the apical cortex of wing and haltere discs. Sum of slices apical intensity projection of L3 wing and haltere cells stained for anti-squash myosin. Myosin shows a polarized accumulation primarily at the apical cortex of both wing and haltere discs. However, this accumulation is relatively less pronounced in haltere cells compared to wing cells, Scale bar,10 μm. C. Quantification of apical myosin mean fluorescence intensity normalised with the wing at L3. The mean and s.d are presented, *** p<0.001, Student’s t-test, n=11 discs for both wing and haltere. D. Wing and haltere disc cells are shown before and after the laser ablation, red dots denote the ablated cell bonds Note the vertex displacement before (t=0 s) and after (t=0.65 s) the ablation (left). Scale bar, 2 μm. E. Quantification of initial mean displacement and F. initial recoil velocities normalised with haltere are plotted. n=17 and 20 cuts for wing and haltere disc, respectively. The mean and s.d are presented, **** p<0.0001, Mann-Whitney test.

Smaller apical surface area and higher levels of actin and Myosin II at the apical cortex of wing disc cells suggest more pronounced apical constriction in wing disc cells compared to those of haltere discs at L3 (A. C. Martin & Goldstein, 2014). This could also be the reason for differences in the cell height (wing disc cells are longer) between the two discs.

### Haltere disc cells have lower apical junctional contractility compared to wing disc cells

Apico-cortical actomyosin accumulation is also correlated with higher cellular tension/contractility apart from apical constriction (Fernandez-Gonzalez et al., 2009; Heisenberg & Bellaïche, 2013; Rauzi et al., 2010). We, therefore, investigated potential differences in the cell apical junctional tension between wing and haltere cells at L3. To achieve this, we conducted laser ablation on the apical pouch region of the wing and haltere disc cells expressing *endo-DE cadherin* GFP. Freshly dissected wing and haltere discs were cultured ex-vivo during the entire duration of the experiment (see materials and methods). A high-power two-photon laser (800nm) was used to physically break the apical membrane of disc cells. The recoil or vertex displacement thus measured is a relative indicator of the cell tension under the same experimental conditions (Farhadifar et al., 2007; Sui et al., 2018). Our results showed significantly lower vertex displacement and initial recoil velocity in haltere disc cells (measured after 0.65 s) in comparison to wing cells, indicating a higher contractility in wing discs (Fig 2D, E, F, movie 1-2).

Taken together, we report that wing disc cells are laterally more elongated compared to haltere disc cells. Wing disc cells have higher levels of actin and myosin at the apical cortex, presumably causing the observed apical constriction and higher degree of apical cell contractility. Our findings indicate a previously unreported distinctions in the morphological and mechanical properties of wing and haltere DP cells, despite both being composed of pseudostratified columnar cells of common origin.

### Ubx modulates cell size and shape in the developing haltere

The presence of Ultrabithorax (Ubx) is vital and sufficient to confer haltere identity by modifying the default wing fate (Cabrera et al., 1985; Castelli-Gair et al., 1990; Lewis, 1978; White & Akam, 1985; White & Wilcox, 1985). We have observed distinct differences between wing and haltere disc cells. RNAi-mediated knock-down of Ubx using *Ubx*-GAL4 resulted in haltere-to-wing transformations, although resulted in flat ‘winglets” (Fig 3A). Individual DP cells were more elongated and apically constricted, resembling the cellular characteristics observed in the wild-type wing disc at L3 (Fig 4A, B, C, D green). The haltere imaginal discs of *Ubx^RNAi^* were bigger compared to the control discs, suggesting an increase in cell number too. They also exhibited higher actomyosin levels and enhanced cell contractility in the apical region of the disc proper (DP) cells (Fig 4B, E, F, G, H, I).

**Figure 3.**
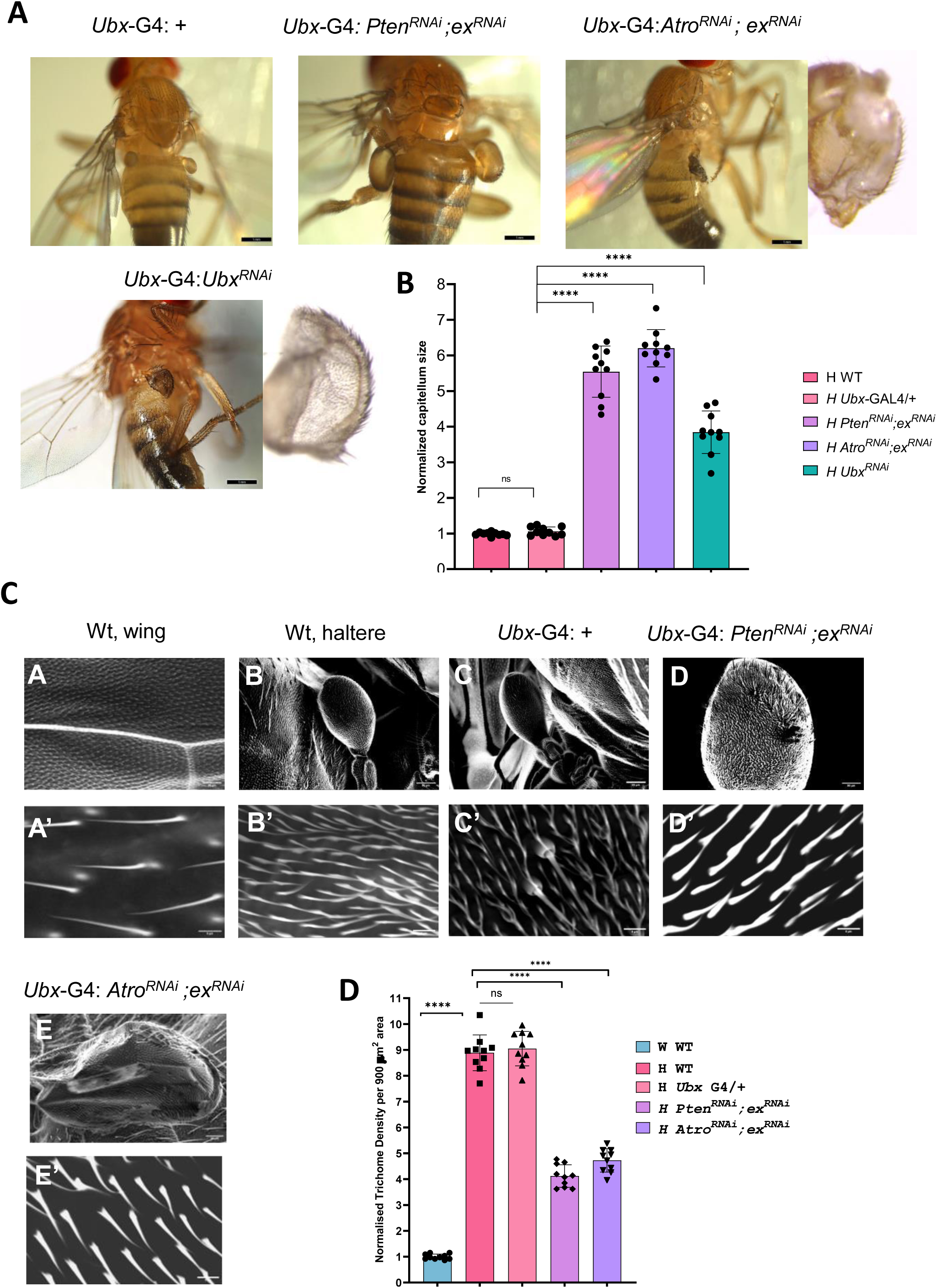
Deregulation of hippo pathway with *Atro* or *Pten* causes partial homeotic transformations. A. Flies with overgrown halteres of indicated genotypes*-Ubx-*G4 (control), *Pten^RNAi^; ex^RNAi^, Atro^RNAi^; ex^RNAi^*and *Ubx^RNAi^*, respectively. *Pten^RNAi^; ex^RNAi^* haltere capitellums were significantly larger, but remained bulbous. *Atro^RNAi^; ex^RNAi^* and *Ubx^RNAi^* flies had halteres which were flatter and showed areas of apposition as well as blistering. Scale bar, 1mm. B. Quantification of haltere capitellum size normalised with the wild-type haltere, the mean and s.d and individual data points are presented, **** p<0.0001, Student’s t-test, n≥10 halteres for each genotype. C. Scanning electron microscope (SEM) images of halteres of the indicated genotypes, except A&A’, which is a representative wing. The top panel shows the overall size and morphology of the organs. A higher magnification of representative area is shown in the bottom panel to visualise the trichome density and arrangement. Note that the (E) *Atro^RNAi^; ex^RNAi^* halteres were larger and flatter. Scale bars: 50 and 5 μm for the upper and lower panels, respectively. D. Quantification of trichome density in a 30 X 30 µm^2^ area, normalised to the trichome density in the wings. The mean, s.d and individual data points are presented, **** p<0.0001, Student’s t-test, n=10 halteres for each genotype.

**Figure 4.**
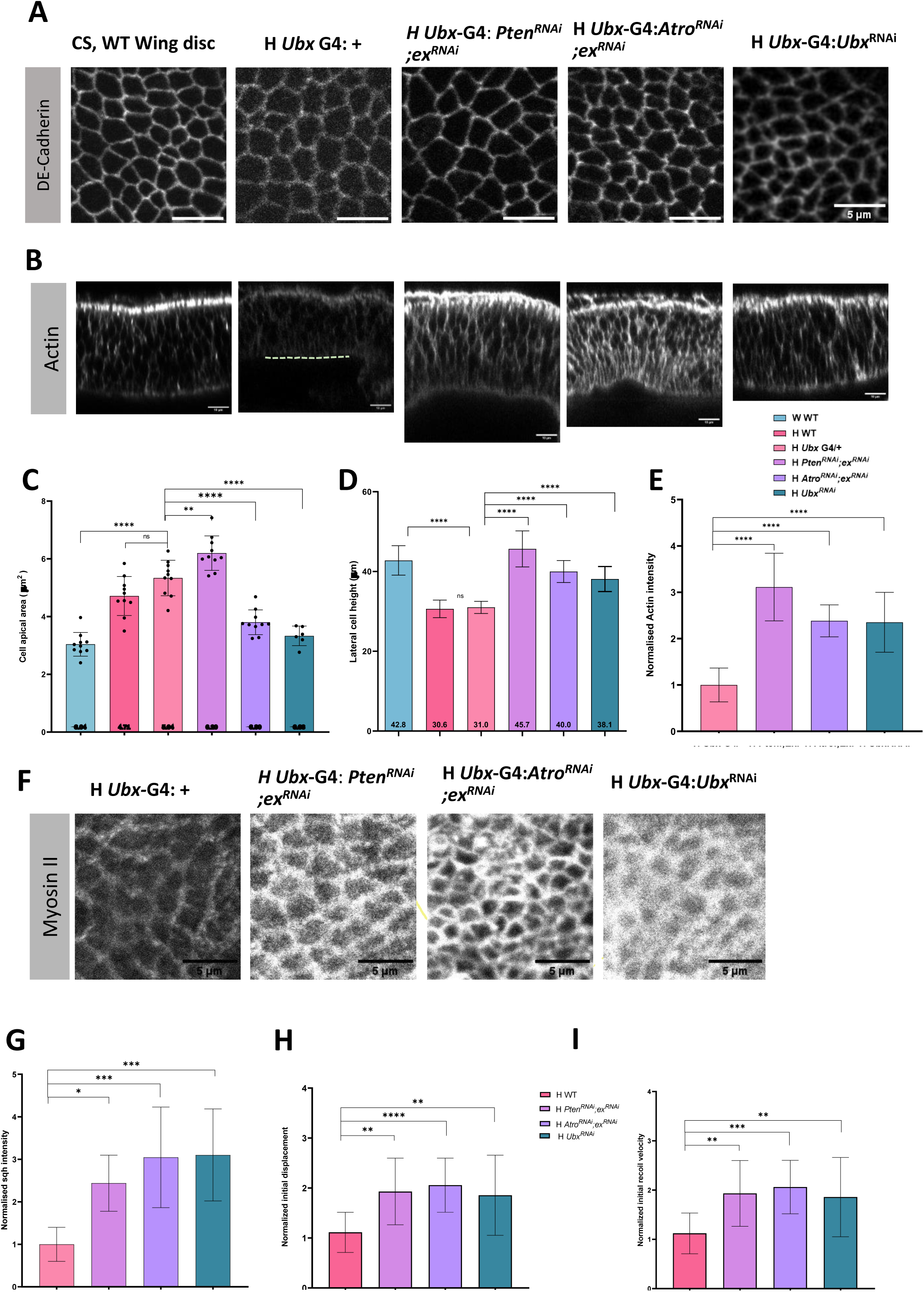
Effect of *Atro^RNAi^; ex^RNAi^* and *Pten^RNAi^; ex^RNAi^* on cellular morphology and cytoskeletal tension in haltere discs during L3 stage. A. Haltere disc proper cells with indicated genotypes expressing DE cadherin to visualise the apical cell membranes. A control wing disc is shown for comparison (left). While *Pten^RNAi^; ex^RNAi^* cells showed a slight increase in the apical cell area, *Atro^RNAi^; ex^RNAi^* and *Ubx^RNAi^* cells showed a reduction comparable to the control wing cells. Scale bar, 5 μm. B. Lateral cross-section of DP cells with genotypes indicated in the above panel stained with actin showing the respective cell height (pouch). Yellow dotted line marks the basal boundary in the *Ubx*-G4 control. *Pten^RNAi^; ex^RNAi^*, *Atro^RNAi^; ex^RNAi^* and *Ubx^RNAi^ -* all shows an elevated lateral cell thickness compared to the control halteres Scale bar, 10 μm. C. Quantification of the cell apical area of mutants with respect to the control wing and haltere discs. Mean, s.d and individual values are represented, ** p<0.01, *** p<0.001, **** p<0.0001, one-way ANOVA, n=10 discs for each genotype except for *Ubx^RNAi^*(n=6). H denotes haltere. D. Quantification of the lateral cell height of the mutants are shown in the plot. n≥7 discs, mean value is shown in the bottom of the bars, mean and s.d are presented, **** p<0.0001, one-way ANOVA. E. Quantification of mean Actin fluorescence intensity, normalised with control halteres. n≥7, mean and s.d are presented, **** p<0.0001, one-way ANOVA. F. Sum of slices-apical intensity projection of haltere disc cells of the indicated genotypes stained for myosin at L3. *Pten^RNAi^; ex^RNAi^*, *Atro^RNAi^; ex^RNAi^* and *Ubx^RNAi^* showed an increased myosin accumulation at the apical cell cortex Scale bar, 5 μm. G. Quantification of Sqh-myosin II mean fluorescence intensity, normalised with *Ubx-*GAL4 control haltere. Mean and s.d are presented, n=9,8,7,10 respectively, * p<0.05 *** p<0.001, one-way ANOVA. H, I Quantification of displacement (H) and recoil velocities (I) normalised with control haltere disc after 0.65 s of ablation. Mean and s.d are presented, n= 17,18,20,19 cuts, * p<0.05, ** p<0.01, *** p<0.001, **** p<0.0001, one-way ANOVA.

Overall, our findings indicate a shift in cellular characteristics from haltere-like to wing-like when *Ubx* is downregulated in haltere cells. While Ubx’s role in regulating organ size via cell proliferation is established (Crickmore & Mann, 2006; Makhijani et al., 2007), our study provides direct evidence of its influence in shaping the cellular three-dimensional morphology and thereby, modulating its mechanical properties. This suggests that Ubx-mediated specification of haltere fate involves modulation of cellular mechanical properties.

### Modulation of haltere growth by the Hippo pathway

The Hippo pathway is one of the critical targets of Ubx (Singh et al., 2015). The Hippo pathway is known to respond to and communicate the cellular mechanical cues through its upstream kinases and the transcriptional activator Yorkie/Yki (YAP in mammals) to regulate growth (Misra & Irvine, 2018; Pan, 2010; Zheng & Pan, 2019). Moreover, it cooperates with many other signalling and patterning pathways, such as EGFR, Insulin and Wnt/Wg to regulate growth and tissue architecture (Wittkorn et al., 2015; Gokhale & Shingleton, 2015; Hansen et al., 2015). Ubx regulates many of the Hippo pathway components and thus, they are differentially expressed between wing and haltere discs (Singh et al., 2015). We examined if differences in the shape, size and mechanical properties between wing and haltere disc cells are due to modulation of Yki pathway by Ubx. Over-expression of Yki did not alter haltere cell shape or size (Huang et al., 2005; Singh et al., 2015). Our group has previously reported that RNAi-mediated downregulation of certain genes in the background of elevated Yki causes massive outgrowth of wing discs, often leading to neoplastic transformations (Groth et al., 2020; Nagarkar et al., 2020). We decided to re-examine those genes for haltere phenotypes. We had earlier reported that RNAi-mediated downregulation of *expanded (ex),* also a component of hippo pathway which negatively regulates Yki mediated growth, gives stronger phenotypes both at larval and adult stages compared to over-expressing *Yki* (Singh et al., 2015).

We, therefore, re-screened previously identified potential negative regulators of growth in the background of reduced *ex* (using the *Ubx*-GAL4 driver) to identify genetic backgrounds that may cause haltere to wing homeotic transformations, specifically flat organ structure, increased organ size and sparse trichome arrangement.

We observed that RNAi-mediated downregulation of *Atrophin* (*Atro*, also known as *Grunge*) or *Pten*, in the background of reduced *ex* resulted in significantly overgrown halteres in adult flies. Atrophin is a transcriptional co-repressor involved in multiple developmental processes including tissue differentiation and patterning (Charroux et al., 2006; Erkner et al., 2002; Wang et al., 2006, 2008; Wang & Tsai, 2008; Yeung et al., 2017). Pten suppresses growth mediated by the insulin pathway (Akt) (Chen et al., 2018; Gao et al., 2000; Goberdhan et al., 1999). Mutants of Pten often display increased cell size and defects in Myosin II localization (Bardet et al., 2013; Goberdhan et al., 1999). Being a Phosphatase, Pten influences the membrane localisation and activation of numerous other proteins involved in growth regulation.

We observed that *Pten^RNAi^; ex^RNAi^* adult flies exhibited larger but bulbous halteres with wing like marginal bristles whereas flies with *Atro^RNAi^; ex^RNAi^*, displayed larger, flatter-“wing-like” halteres also with marginal bristles (Fig 3A, C). To estimate the increase in organ size, we quantified the size of haltere capitellum relative to the control haltere (Fig 3B). While, there was no significant difference between the wildtype and *Ubx*-GAL4 control (without any UAS transgene) capitellum size, the *Atro^RNAi^; ex^RNAi^* and *Pten^RNAi^; ex^RNAi^* halteres were at least 5 folds larger compared to the controls. In addition, the halteres from *Atro^RNAi^; ex^RNAi^* flies also showed areas of partial apposition and small blistered phenotype (Fig 3A). For comparison, adult haltere phenotypes of *Ubx^RNAi^*flies are shown in the same panel.

The arrangement of trichomes and its density can serve as an indirect indicator of cell size and polarity (Singh et al., 2015). In the adult wing, each cell typically bears only one trichome, and the cells are larger hence, the trichomes appear to be sparsely arranged on the cell surface. In contrast, haltere cells are smaller, and they may have more than one trichome per cell. As a result, the trichomes on haltere cells appear denser (Roch & Akam, 2000). Scanning electron micrographs revealed that the arrangement of trichomes in *Atro^RNAi^; ex^RNAi^* or *Pten^RNAi^; ex^RNAi^*halteres, was sparser when compared to control halteres (Fig 3C). To further test this, we did a quantitative analysis of trichome density by counting the number of trichomes within a defined area at various locations on the wing blade as well as on the haltere capitellum of different genotypes. A noticeable decrease in trichome density, upto 50%, was observed in both *Atro^RNAi^; ex^RNAi^*and *Pten^RNAi^; ex^RNAi^* flies (Fig 3D). The sparser arrangement of trichomes in the *Atro^RNAi^; ex^RNAi^*and *Pten^RNAi^; ex^RNAi^* halteres is an indication of increased cell size and flattening.

Hippo pathway regulators such as Yki and ex are known to regulate growth via increasing cell proliferation, thereby the cell number (Irvine & Harvey, 2015, Singh et al., 2015). Pten downregulation is known to increase cell growth and cell area (Gao et al., 2000). Hence, the pronounced phenotypes observed in *Atro^RNAi^; ex^RNAi^* or *Pten^RNAi^; ex^RNAi^* flies may potentially be due to breakdown of the constraints caused by strong inverse correlation between cell size and cell number during organogenesis.

### *Atro^RNAi^; ex^RNAi^* and *Pten^RNAi^; ex^RNAi^* haltere disc cells show increased cell height, apical cell constriction and changes in cellular morphology

In order to understand the effect of *Atro^RNAi^; ex^RNAi^*and *Pten^RNAi^; ex^RNAi^* on cell morphology, we measured cellular apical area and lateral height of haltere disc cells at L3 stages. Interestingly, while the haltere disc cells of the *Atro^RNAi^; ex^RNAi^* showed a decrease in the cell apical area compared to the control haltere disc cells, *Pten^RNAi^; ex^RNAi^* showed an increase in the apical cell surface area (Fig 4A, C).

Next, we measured the lateral cell height of the DP cells in these mutant haltere discs. We observed an increase in the cell height of both *Atro^RNAi^; ex^RNAi^* and *Pten^RNAi^; ex^RNAi^* haltere disc cells (Fig 4B, D). The cells were significantly elongated compared to those in wild type haltere discs and were comparable to cells of the wild-type wing disc. Some *Pten^RNAi^; ex^RNAi^* haltere cells were approximately 55 microns in length, much longer than what was normally observed for even wild-type wing disc cells.

To examine whether the aforementioned cell shape changes are accompanied by alterations in the actomyosin cytoskeleton, we estimated F-actin and Myosin II levels at the cell junctions in the two mutants. Notably, we found that both F-actin and Myosin II (Sqh) were enriched at the apical surfaces of cells of the mutant haltere discs compared to cells of the control haltere disc at L3 (Fig 4B, E, F, G).

The alterations in cell morphology and the higher levels of actomyosin at the apical belt potentially signify changes in cellular and tissue tension in the mutant halteres. To evaluate this possibility, we measured apical cell contractility in DP cells by laser ablation. We observed that cells of mutant haltere discs exhibit higher initial mean displacement and recoil velocity compared to the wild type discs, indicating an increased tension at the apical membrane of these columnar cells (Fig 4 H, I, movie 2-5, S3).

Taken together, in the light of our earlier observations, these findings indicate that both *Atro^RNAi^; ex^RNAi^* and *Pten^RNAi^; ex^RNAi^* halteres undergo cellular-level homeotic transformations even at L3, albeit to varying degrees. *Atro^RNAi^; ex^RNAi^* haltere disc cells exhibit nearly complete haltere to wing transformation, characterized by enhanced cell elongation, apical constriction, elevated actomyosin levels, and increased tension at cell junctions compared to the wildtype haltere disc at L3. Changes in cell contractility and tension are reported to not only drive two-dimensional shape deformation; they also impact cell size and proliferation (for instance via the hippo pathway;(Dawes-Hoang et al., 2005; Liu et al., 2022; Polyakov et al.,2014) collectively influencing organ size in a three-dimensional context.

### Basal ECM remodelling and DV apposition during haltere morphogenesis

Towards the end of L3, wing and haltere discs begin complex folding to undergo eversion, resulting in the formation of a bilayer structure. In everting wing discs, dorsal-ventral (DV) columnar cells appose each other basally, without much space / lumen inside, while in everting haltere discs have distinct lumen keeping the two layers separate (De las Heras et al., 2018; Diaz-de-la-Loza et al., 2018). Earlier reports suggest that removal of *Ubx* or inducing the expression of *MMP*s, results in the flattening of everting haltere discs (De Las Heras et al., 2018). Conversely, inducing the expression of Timp, which inhibits MMPs or downregulating integrins results in blistering in the adult wings (S4) (De las Heras et al., 2018; Domínguez-Giménez et al., 2007). Interestingly, it was also noted that the remodelling of bECM is more critical compared to the aECM for the DV apposition and organ flattening (Diaz-de-la-Loza et al., 2020).

In line with the previous studies, we also observed a clear bECM remodelling and DV apposition in wild type wing discs at 4-6h APF compared to haltere discs (Fig 5A, B) (De las Heras et al., 2018; Diaz-de-la-Loza et al., 2018). The *Atro^RNAi^; ex^RNAi^* adult halteres were larger and flattened with areas of partial apposition (Fig 3A, C), whereas *Pten^RNAi^; ex^RNAi^* adult halteres are bulbous, although larger.

**Figure 5.**
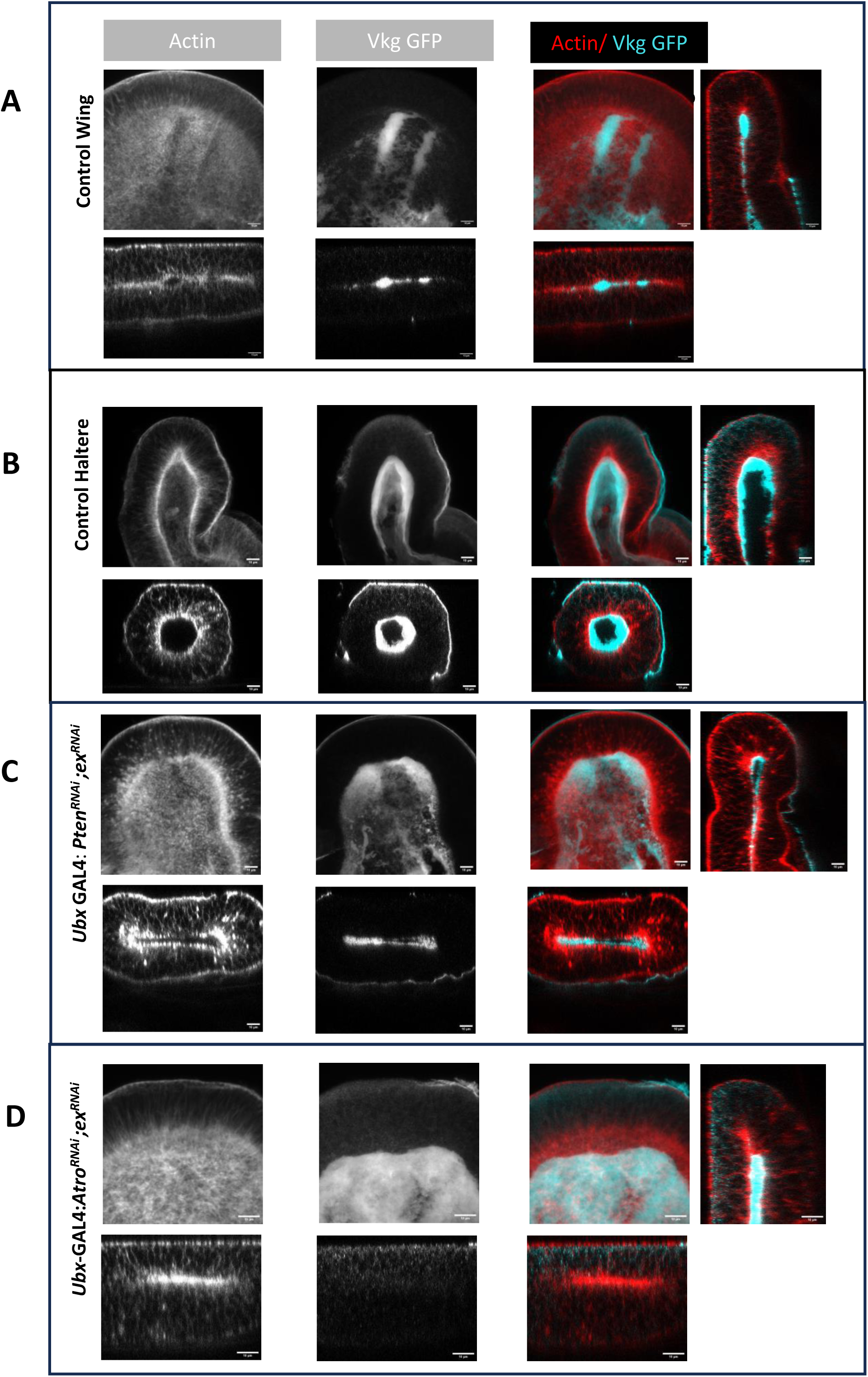
Basal ECM remodelling and 3D tissue shape deformations in partially transformed pupal haltere discs (4-6hAPF). A. *Drosophila* wing (A) and haltere discs (B) stained for F-actin (red) and Vkg GFP (cyan), respectively. Z-projection of the pouch (a sum of slices spanning the bECM) is shown. Individual channels are shown in greyscale, with their respective cross-sections. Note the Vkg GFP accumulation in the lumen of haltere discs compared to the wing discs. Also, the control haltere exhibits a more globular architecture, while wing discs tend to be more elliptical in cross sections. Scale bar, 10 μm. B. *Pten^RNAi^; ex^RNAi^* haltere disc. The cross-sections of the discs appear to be less rounded, indicating a 3D deformation towards a flatter geometry. The presence of Vkg GFP signal was observed throughout the discs. Additionally, in some individuals, there was an increased lumen observed, plausibly making the overall structure more globular. Scale bar, 10 μm. C. *Atro^RNAi^; ex^RNAi^* haltere disc. There is an overall tissue flattening that was observed in *Atro^RNAi^; ex^RNAi^* haltere discs. As in the developing wing, the *Atro^RNAi^; ex^RNAi^* shows areas of ECM degradation– indicated by lack of Vkg GFP signal and tight basal-basal zippering of the dorsal and ventral layers in those areas. Scale bar, 10 μm.

To investigate the status of bECM, we generated fly lines expressing Vkg-GFP with *Atro^RNAi^; ex^RNAi^* and *Pten^RNAi^; ex^RNAi^*. The respective imaginal discs were stained for F-Actin and the bECM marker Vkg. Extreme care was taken to minimise any flattening of the tissue while processing for staining and imaging. Interestingly, at 4-6h APF, we observed *Atro^RNAi^; ex^RNAi^*everting haltere discs with areas of basal ECM degradation and DV apposition, evidenced by the absence of Vkg GFP signal and the lack of a distinct lumen between the DV layers (Fig 5D). *Pten^RNAi^; ex^RNAi^*discs did not show such signs of bECM clearance (Fig 5C). The Vkg signal was consistently detected in all cross-sections and across all the tested discs. The lumen was intact and no DV apposition was observed (Fig 5C). However, an overall 3D deformation and flattening of the tissue geometry was seen in some discs. Thus, probably due to the lack of bECM clearance and the DV apposition, the *Pten^RNAi^; ex^RNAi^*adult halteres retained their bulbous shape, although sparse arrangements of trichomes suggest wing-type flat cells. Thus, the degree of homeotic transformation is likely to be lower in this genetic combination.

### Modelling cell shape transitions and organ morphogenesis

The experimental studies presented here demonstrated that the initial shape difference between the cells of wing and haltere discs (at L3 and at 3-4hAPF) is potentially due to the differences in heights, surface area, and contractility between the individual cells of wing and haltere discs. Moreover, we also find that, starting from this initial state, different experimental conditions involving cell growth and proliferation, ECM degradation, cellular adhesivities and contractilities, etc., lead to different overall organ shape for haltere and wing. However, we lack direct experimental evidence beyond 6-8hAPF, specifically for developing haltere. As a result, how underlying physical factors involved in these cellular and disc level differences ultimately lead to the adult wing and haltere morphology are less understood. We made an attempt to understand the relationship between the observed parameters in the initial and the final shape of the organs using simple theoretical consideration. Specifically, we used lateral section 2D vertex model to qualitatively understand how the interplay between adhesivity at different locations (apical, basal, and lateral), ECM degradation, and lumen dynamics modulate the shape of individual cells and the overall tissue shape.

As shown in Fig 6A, in the lateral vertex model, we represent each of the cells as a quadrilateral with four vertices and four edges. The apical edge of the cell with line tension of *λ*_*a*_ is exposed to the surrounding environment. The bottom or the basal edge of any cell is in contact with the central lumen and has a contractility of *λ*_*b*_. Any cell has two lateral edges with contractility *λ*_*l*_ such that each lateral edge is shared between the neighbouring cells. The cells are forced to maintain the effective area *A*_0*c*_ using quadratic energy penalty. The apical and basal ECM are modelled using linear springs of stiffness *k*_*a*_ and *k*_*b*_, respectively. The rest length of the spring corresponds to the initial length of the corresponding edge. The ECM corresponding to the linear springs resists changes in apical and basal lengths. The central lumen is provided with bulk stiffness of *K_lum_* (Fig 6A) and preferred area *A*_0*l*_which together control the lumen pressure and size. In our model we dynamically modulate *A*_0*l*_ as a surrogate for cytoplasmic deposition from the cells into the lumen or fluid exchanges between the lumen and the exterior. The overall dynamics of these factors dictate the shape of individual cells and the overall organ geometry in our model. More *Mathematical details of the model* are given in *Material and Methods*.

**Figure 6.**
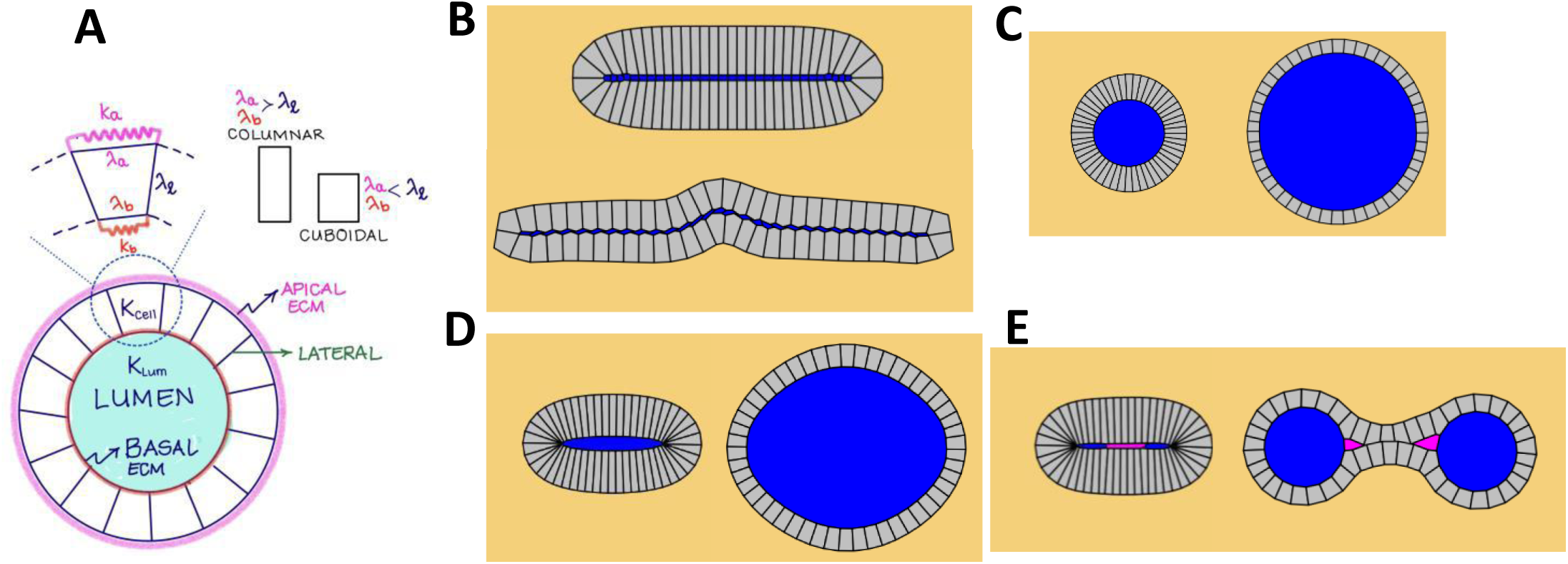
Modelling of cell and organ shape transitions during wing and haltere morphogenesis. **A.** The schematic showing different components of a lateral cut haltere model with a spherical geometry. Various components of the model are discussed in the text. The apical, basal, and lateral contractilities are *λ*_*a*_, *λ*_*b*_, and *λ*_*l*_, respectively. Individual cell has bulk stiffness of *K*_cell_ and preferred area *A*_0*c*_. The apical and basal ECM are represented as linear springs of stiffnesses *k*_*a*_ and *k*_*b*_, respectively. The central lumen is provided a preferred area *A*_0*l*_ and bulk stiffness *K*_lum_. For the columnar cells, *λ*_*a*_, *λ*_*b*_ > *λ*_*l*_, whereas for cuboidal cells we have *λ*_*l*_ > *λ*_*b*_, *λ*_*a*_. **B.** At the 4-6h APF, the wing has two apposed layers of columnar cells that are adhered to each other by a layer of integrins (blue layer of cells). A reduction in apico-basal contractilities and increase in lateral contractility along with ECM degradation results in the columnar to cuboidal cell transition. Because of the lack of central lumen and adhesion between the top and bottom cell layers, unlike haltere, the wing retains flat morphology. C. For WT haltere, the initial condition corresponds to columnar cells surrounding a central lumen (blue). A reduction in apico-basal contractilities and increase in lateral contractility while accompanied with a continuous lumen increase and ECM degradation results in the maintenance of the final globular haltere shape (right) that is now lined with cuboidal cells. D. In the *Pten^RNAi^; ex^RNAi^* haltere mutant, the initial condition is very similar to that of the wing. However, the top and the bottom layers do not adhere to each other and there is a small central lumen. Consequently, any increase in lumen size due to material exchange ultimately results in a globular haltere. As in the WT haltere case, the final geometry of the individual cells can undergo a columnar to cuboidal transition if modifications in contractilities happen. E. The *Atro^RNAi^; ex^RNAi^*haltere mutant, as the *Pten^RNAi^; ex^RNAi^* mutant, has a wing-like flat initial arrangement of columnar cells. However, the top and the bottom layers are partly adhered to each other (pink connection) while encompassing a small lumen (blue). The adhesion of the central layer in this case, while the growth of the lumen results in a final shape that is partly adhered like wing while partly being globular like haltere.

**Figure 7.**
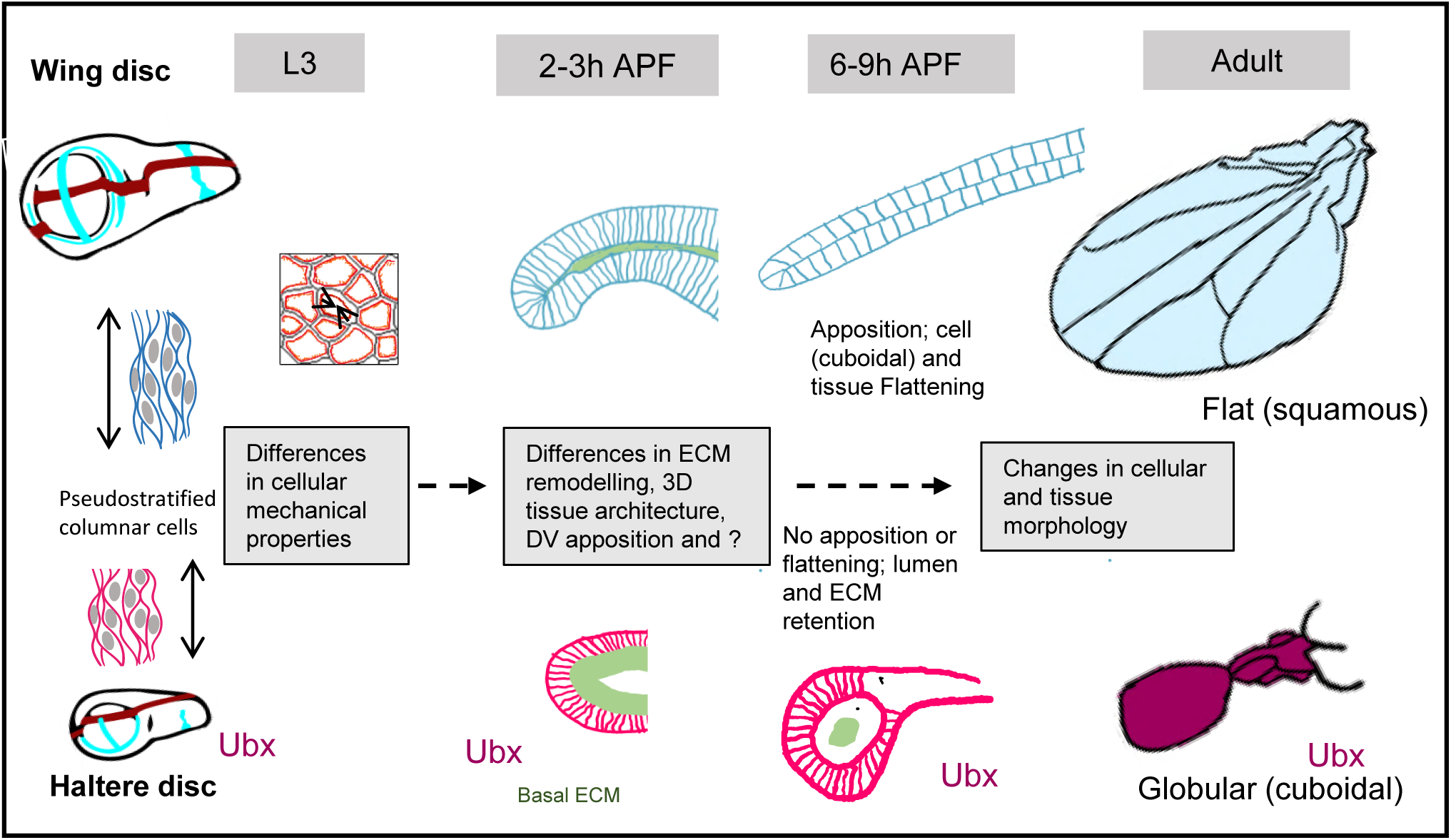
Schematic showing the summary of events during the wing and haltere morphogenesis.

### Wild type Wing

We started with the wing-disc geometry at 4-6APF in which two layers of columnar cells are apposed and zippered to each other without any lumen (Figure 5A, 6B top). As reported earlier, the ECM degradation at the basal layer of the wing is important for the zippering of these two layers (De las Heras et al., 2018; Diaz-de-la-Loza et al., 2018, 2020). When *λ*_*a*_, *λ*_*b*_ > *λ*_*l*_, i.e., when the lateral contractility of the cells is smaller when compared with their basal and apical counterparts, each of the individual cells will be columnar, i.e., their lateral dimensions would be larger when compared to their apical and basal sizes (Fig 6B, top). When, due to localization of myosin to the lateral side (as reported in Diaz-de-la-Loza et al., 2018), the lateral contractility becomes gradually larger (Eq. 3) as compared to the apical and basal side, i.e., *λ*_*l*_ > *λ*_*a*_, *λ*_*b*_, the cells change their shape from columnar to cuboidal. This happens when the apical and basal ECM is simultaneously degraded (*k*_*a*_ = *k*_*b*_ = 0), thus facilitating changes in individual cell morphologies (Eq. 4). However, since the cells in the top and the bottom layer (or D and V layers) are connected to each other with the help of integrins (Domínguez-Giménez et al., 2007; T. Sun et al., 2021), this leads to flattening and expansion of the overall wing geometry along with the transformation of the individual cells from columnar to cuboidal (Fig 6B, right).

### Wildtype haltere

Our observations along with previously published reports suggest that the cross-section of haltere pouch at 4-6hAPF has a globular overall geometry (Fig 5B) (De las Heras et al., 2018). Thus, we modelled haltere with an initial globular geometry as depicted in Fig 6C. Although there is relatively less actin-myosin in the apical cortex and apical tension, qualitatively, the observed columnar shape of individual haltere cells at larval stages would still be due to *λ*_*a*_, *λ*_*b*_ > *λ*_*l*_. As reported earlier and observed here, at 4-6h APF, in the absence of ECM degradation, the dorsal and ventral layers are not connected to each other and thereby leading to the formation of pronounced lumen (De las Heras et al., 2018). This makes the morphogenesis of the two layers decoupled from each other.

The presence of lumen provides an initial globular geometry to the haltere. However, for such geometries, the overall shape of the tissue is strongly coupled with the basal and apical areas of individual cells. Consequently, the size of lumen and morphology of individual cells are related to each other. In our simulations, the contractilities are gradually modified (Eq. 3) to reach values that are compatible with the columnar geometry of individual cells (*λ*_*l*_ > *λ*_*a*_, *λ*_*b*_). Simultaneously, we also introduced delayed decay (Eq. 4) of ECM (as observed experimentally) and a similar slow pace in the increase in the lumen size. Altogether, we observed a gradual transition of the individual cell morphology from columnar to cuboidal and a corresponding increase in haltere size, while it retains spherical shape as is observed experimentally. The dynamical evolution of the model haltere from its initial morphology to the final organ shape is shown in Fig 6C. On the other hand, if the individual cells undergo columnar to cuboidal shape change prematurely before the lumen is sufficiently large, the overall haltere geometry becomes temporarily wavy to accommodate the additional basal area. We note that the technical measures implemented in the haltere simulation to ensure the absence of this waviness, i.e., using longer time scales (*t*_0*s*_, *t*_0_, τ, τ_*s*_, Eqs. 2 and 3) for gradual modifications of line tensions and ECM degradation (apical and basal springs) as compared to that for the wing (Table 1), result in a larger time to achieve the final haltere geometry (Fig 6C) when compared with the wing counterpart (Fig 6B). This is consistent with the observed developmental time between wing and the haltere cells to undergo columnar to cuboidal transitions.

### Pten^RNAi^; ex^RNAi^ haltere

*Pten^RNAi^; ex^RNAi^* larval haltere cells are similar to wing cells in lateral height, the levels of apical acto-myosin complex and the apical contractility (Fig 4). During 4-6h APF, although we did not observe ECM degradation, the cross-section of discs were flatter compared to the wildtype haltere discs (Fig 5B, C). However, there was a presence of small lumen keeping the dorsal and the ventral layers separate from each other.

Interestingly, the outcome of the simulation for these mutant discs was a geometry very similar to that of wildtype haltere discs (Fig 6D). *Pten^RNAi^; ex^RNAi^*adult halteres too showed overall globular morphology (Fig 3A, C). This suggests due to the lack of coupling between the dorsal and the ventral layer, increase in the size of the lumen is not resisted resulting in the overall globular geometry.

### Atro^RNAi^; ex^RNAi^ haltere

*Atro^RNAi^; ex^RNAi^* larval haltere cells are also similar to wing cells in lateral height, the levels of apical acto-myosin complex, apical constriction and the apical contractility (Fig 4). During early pupal stages (4-6hAPF), the discs were flatter compared wildtype haltere discs (Fig 5B, D). However, unlike *Pten^RNAi^; ex^RNAi^*, we did observe ECM degradation. The dorsal and the ventral layers were closely opposed to each other, although heterogeneously (Fig 5D).

With this starting point, the outcome of the simulation for these mutant discs was flat wing-like geometry at the locations wherever the dorsal and the ventral layers are coupled (Fig 6E, adhesive pink region). In other locations (Fig 6E, blue growing lumen), wherever this coupling was absent, we observed globular geometry giving an overall blistered appearance as observed experimentally (Fig 6E, right).

Taken together, our results reported here suggest that major differences between wing and haltere development are set in late L3 stages, specifically in lateral height and apical constriction and contractility. These differences coupled with delayed degradation of ECM in haltere discs provides clues on how distinct morphological shapes that we observe in adult wings and halteres arise.

## DISCUSSION

While it is obvious that shape and size of tissues and organs are dictated by the properties of individual cells, it is not clear what properties of cells dictate how much and which aspects of organ size and shape. In this study, we present novel findings regarding differences in cellular characteristics, including size, shape, and contractility, between the third instar wing and haltere disc cells, which are key for those organs to attain specific size and shape. At L3, wing and haltere discs are made up of pseudostratified columnar cells. A detailed examination of these discs during the third instar larval stage revealed that wing cells were more laterally elongated with a narrower apical surface compared to haltere cells. Additionally, we observed higher levels of apical actin and myosin II, key regulators of cell tension and contractility, in wing disc cells. Quantification of apical contractility of wing and haltere disc cells using laser ablation confirmed higher degree of contractility in wing disc cells as compared to haltere disc cells. Cell sorting observed in *Ubx^-^*clones from its surrounding wildtype cells (Ubx^+^) in halteres suggest that, in addition to the differences in cell size, shape, apical Actomyosin accumulation and contractility reported in this study, there may also be differential cell adhesion properties between wing and haltere cells (Morata and Garcia-Bellido, 1976; Shashidhara et al., 1999).

Reducing the Ubx levels using *Ubx^RNAi^* in haltere discs resulted in an increase in the actomyosin levels at the apical cortex. The cells were apically constricted and laterally elongated, contrasting with the wildtype halteres. This observation indicated a transformation in cell characteristics towards a more wing-like phenotype. The resultant adult morphology was flatter ‘winglets’ in place of globular halteres, albeit smaller in size (compared to wings), with a sparser trichome arrangement. This suggests a previously unrecognised role of Ubx in altering the cellular mechanical properties of haltere cells as early as the third instar larval stage.

Wing and haltere disc cells, like other epithelial cells, display apicobasal polarity, and the mechanical properties of these cells differ across their intracellular domains. Modifying the properties of these domains has the potential to alter cell shape and, consequently, the overall tissue architecture. For instance, at the subcellular level, differential actomyosin accumulation-induced constriction drives tissue bending (Heer & Martin, 2017; Heisenberg & Bellaïche, 2013). Furthermore, cells are mechanically coupled to their neighbours through adherent junctions, coordinating the forces generated by individual cells to be integrated across the tissue. Our simple vertex model suggests that the transition between columnar (elongated) and cuboidal cell shapes can be facilitated by adjusting cell contractilities. Specifically, modifying parameters such as apical, basal and lateral cell contractility allows us to shift cell shapes from a columnar to a cuboidal morphology and vice versa. Cuboidal cell shapes emerge when lateral cell contractility exceeds both apical and basal contractility, and this process involves tissue deformation, in this case, flattening.

Additionally, cells are surrounded by ECM at apical and basal surfaces, which also acts as a controllable mechanical constraint for morphogenesis (Harmansa et al., 2023). Previous studies have shown that wing discs undergo columnar to cuboidal cell shape at early pupal stages, which correlates with ECM remodelling. Halteres, on the other hand, do not undergo cell shape transition or ECM remodelling. A compromised Ubx level in halteres using *Ubx^RNAi^*, induces ECM degradation and an overall flatter geometry (De las Heras et al., 2018; Diaz-de-la-Loza et al., 2018, 2020). Incorporating ECM in our computational model predicts that, clearance of ECM is necessary to change from columnar to cuboidal cell shape.

Taken together, our results suggest that the dynamics of ECM and actomyosin complexes are important in determining the cell and tissue architecture. The alterations in these can have a profound impact on organ development. Hence, the signalling pathways that activate actomyosin contractility, such as hippo and, ECM, are fine-tuned by a balance of both positive and negative regulation by Ubx.

We also identified additional mutant combinations that exhibited varying degrees of haltere-to-wing transformation. In the *Atro^RNAi^; ex^RNAi^* halteres, during the larval stage (L3), we observed that the cells had transitioned towards a more wing-like appearance, exhibiting increased apical constriction, cell height and actomyosin levels, as well as cell tension. At 4-6 APF, we observe varying degrees of heterogeneous basal extracellular matrix (ECM) degradation. Remarkably, in the regions where ECM clearance occurred, we observed tight zippering of the dorsal and ventral layers, alongside an overall flattening of tissue geometry. The *Atro^RNAi^; ex^RNAi^* halteres, after eclosion, displayed areas of partial apposition and small blistered phenotypes. The haltere capitellum was significantly enlarged, with a flatter morphology, resembling “wing-like” (but, much smaller than wildtype wing), complete with marginal bristles and a reduced trichome density.

*Pten^RNAi^; ex^RNAi^* flies also exhibited enlarged halteres, with the presence of wing marginal bristles and a sparser arrangement of trichomes but, retained the bulbous shape. We noted an overall increase in the apical cell size within the *Pten^RNAi^; ex^RNAi^*haltere discs. The cells were elongated with increased levels of actomyosin complexes at the cell cortex, indicative of enhanced cell contractility. This increased contractility was subsequently confirmed through contractility experiments. During the early pupal stages (4-6 APF, we monitored), clearance of the ECM was not observed, which resulted in a distinct lumen with separating D and V layers. Nevertheless, there was an overall 3D shape deformation of the tissue towards a flatter geometry.

The shape of the haltere is highly resistant to changes, as evidenced by our screen and previous literature (Giraud et al., 2021). Apart from Ubx, only a limited number of genes or combinations of genes have been identified with the potential to induce a flatter morphology in the haltere. In the case of the *Pten^RNAi^; ex^RNAi^* mutant, despite undergoing a partial homeotic transformation at the cellular level during the larval and adult stages and exhibiting a somewhat flat geometry at early APF, the adult organ remained globular shape. Evidenced from our computational model, a plausible explanation is that due to the absence of basal ECM degradation and dorsoventral zippering, the lumen was retained, eventually filling with hemolymph, resulting in the adult globular geometry. This observation also implies that the flattening observed in the wings cannot solely be attributed to differences in cell numbers between the wing and haltere. Even if the disc is significantly larger in size (higher cell number), it can still exhibit a globular morphology, highlighting the importance of other factors influencing tissue shape beyond cell numbers.

An interesting enigma arises when considering the influence of extracellular matrix (ECM) degradation on wing and haltere shape generation. In the case of wing discs, preventing basal ECM degradation is sufficient to hinder the apposition of dorsal and ventral layers and render the organ globular. Conversely, inducing ECM degradation by overexpressing MMP1 or reducing Timp expression does not suffice to flatten the halteres completely. For instance, when MMP1 expression is induced in halteres during the mid-late larval stage, it clears the basal ECM and alters individual cell morphology to a cuboidal shape. However, The D and V layers remain separated with an increased lumen size, resulting in adult globular halteres with no apposition or expansion (De las Heras et al., 2018). This uncouples ECM degradation and cell shape transition from dorsal-ventral zippering and 3D flattening of tissue. This, again, emphasises the multi-level control of haltere fate by Ubx, rendering the halteres highly resistant to shape changes, unlike the wing.

Similarly, reducing Ubx levels in halteres using *Ubx^RNAi^*leads to the de-repression of ECM (Vkg) degradation, resulting in flatter cells and tissue, ultimately giving rise to an adult organ with a flat ‘winglet’ morphology. However, the timing of Ubx expression appears to correlate with appendage development and attaining its characteristic shape. Early downregulation of Ubx results in large, flat appendages (small wings), while late downregulation (late L3 onwards) leads to smaller appendages that are globular in shape (De las Heras et al., 2018). This suggests that organ flattening is observed only when MMP1 expression is induced during the early L3 stage, which may cause ECM degradation at very early stages of the pupal development. Interestingly, eliminating integrin also induced a premature shortening of the cell height in the L3 wing discs, which subsequently developed into a blistered wing (Domínguez-Giménez et al., 2007). The authors also note that the timing of the induction plays a crucial role in obtaining the desired phenotype. The integrins levels need to be reduced substantially at least by prepupal stage to give sufficient blistered flies. These observations emphasise that modifications to the halteres must be initiated at least by mid to late-larval stages to substantially impact the organ’s shape. This further supports our findings that the differences in cellular mechanical characteristics between late L3 wing and haltere discs may represent critical factors determining the shape of those organs in adult flies.

## Acknowledgement

We thank Bloomington, Vienna Drosophila Stock centres and Richa Rikhy for fly lines and reagents, Vijay Vithal for help in microscopy, Richa Rikhy, Girish Deshpande, Girish Ratnaparkhi and LSS lab members for their comments and support. This is work is funded by CSIR (to CD), Science and Engineering Research Board (MTR/2020/000605; to MMI) and IISER Pune, Ashoka University and NCBS (to LSS).

## Materials and methods

### Fly stocks and maintenance

The fly stocks for crosses were grown on standard cornmeal agar media and were maintained at 25°C. The wild-type strain used during the study is Canton-S. All the crosses were set up at a temperature of 25°C, unless stated otherwise.

Fly lines used in this study: CS, *vkg*-GFP (BL 98343), Sqh-GFP (BL57145), *endo-DE-cadherin-GFP* (BL60584), *Ubx* GAL4 (Pallavi & Shashidhara, 2003), *Ubx* GAL4^M1^ (de Navas et al., 2006), w*, *ap*-GAL4, UAS-GFP/CyO; UAS-Yki, tub-Gal80ts/TM6B (Song et al., 2017), *nub* GAL4 (from Chaudhary V), UAS *Ubx^RNAi^* (v37823), UAS *Atro^RNAi^* (KK107413, BL51414), UAS *ex^RNAi^* (TRiP.JF03120, KK109281), UAS *Pten^RNAi^* (KK101475), *If ^RNAi^* (BL27544), *Timp^RNAi^* (BL61294).

### Immunohistochemistry

#### Staining of larval discs

Wandering third instar larvae were collected and washed with Phosphate buffer saline (PBS, pH7.2 Invitrogen) in a glass cavity block. After giving a posterior cut to remove maximum fat bodies, the anterior part, which contains the discs of interest, was turned inside out. At least 8 such heads were fixed for 20 minutes in 4% Paraformaldehyde. Fixed larval heads were given 2 washes of 0.1% PBTX for 10min each and were blocked with 0.5% BSA for 2hr at room temperature. Larval heads were then incubated with primary antibodies of the desired dilution overnight at 4°C. This was then followed by two washes for 10 min each with 0.1% PBTX and incubation with the secondary antibody (1:1000) in blocking solution for 2 hr at room temperature in the dark. Then larvae were again given two washes of 10 min each with PBTX. A final wash with PBS was given to remove all PBTX. Imaginal discs of interest were obtained by dissecting them from anterior heads in microscopic slides. Imaginal discs were then mounted with anti-fade mountant (Invitrogen) or Vectorshield and double-sided tape as spacers. All the imaging was done Leica Sp8 confocal microscope using 63X/1.40NA Oil with a magnification of 0.75 or 2.5.

#### Reagents

Phosphate buffer saline PBS pH7.4 Sigma Blocking solution: PBS + 0.1% TritonX-100 (sigma) + 0.5% BSA (Sigma). PBTX: PBS + 0.1% TritonX-100 (sigma).

#### Antibodies Used

Primary antibodies used in this study are anti-N-terminal Ubx 1:1000 dilution (Agrawal et al., 2011), rabbit anti-Sqh (1:400) (Vasquez et al., 2014), rat anti-DE Cadherin (DSHB) (1:50), Rhodamine Phalloidin (1:200) Invitrogen, Rabbit anti-GFP (1:750) Invitrogen, rabbit anti-Viking antibody (1:400 a gift from Stéphane Noselli). All secondary antibody used in this study-Alexa-fluor 568,594,488 and 633 (1:1000) were obtained from Invitrogen.

### Pupal disc ex-vivo culturing

Schneider’s medium from Gibco Thermo Fisher 21720024 was supplemented with 15% FBS and 1% penicillin-streptomycin (15140–122; Invitrogen). Ecdysone from Sigma, 20-hydroxyecdysone H5142, was added at a final concentration of 0.5 μg/mL for attached discs. 0hr or 3hr pupae were washed in sterile PBS and disinfected in 70% ethanol again followed by a wash with 1X PBS. After rinsing in the eversion media, the Pupal case was cut opened in the chilled eversion media. The wing and haltere imaginal discs attached to the pupal walls were then cultured in 48 well dishes at 25°C. After the completion of developmental time of interest, the tissue was fixed in 4% PFA for 20 minutes, and Immunostaining was performed. (Aldaz et al., 2010; De las Heras et al., 2018; Diaz-de-la-Loza et al., 2018).

### Laser ablation experiment

DE-cad-GFP expressing wing and haltere discs of appropriate genotypes were used to visualise cell membranes. The respective discs were carefully dissected in the filter-sterilized chilled Schneider’s media to keep them alive. The discs were mounted in coverslip bottom chambers or on normal slides with appropriate spacers with apical surface facing the bottom coverslip in the same media. Laser ablation was performed using a Zeiss LSM780 microscope equipped with an 800 nm multiphoton femtosecond pulsed Mai Tai laser under a 63X/1.4 oil immersion objective (3-zoom). A reduced area of 13x13 microns was used for the acquisition, and a linear horizontal region of interest (ROI) of approximately 2.5-3.8 microns spanning one cell was ablated using laser power between 45-50%. Imaging was carried out for at least 50 frames after ablations, with a frame duration of 0.57 seconds, except for the frame immediately following an ablation event, which had a duration of 0.64 seconds (Sui et al., 2018). Care was taken to only ablate the ROI and not to damage the cells.

### Quantification of laser ablation

The vertices of the ablated cell edges in the recorded images were tracked manually using ImageJ. To determine the initial mean displacement, we measured the increase in vertex distance between the time point just before ablation (t=0s) and the first image acquired 0.64 seconds after ablation. The initial recoil velocity was calculated by dividing the initial displacement by 0.64 seconds (Sui et al., 2018). This average recoil velocity serves as an indicator of the relative mechanical tension on the cell edge before ablation.

### Cell height

Respective imaginal discs stained with phalloidin and DAPI were mounted carefully on coverslips without flattening. The images were acquired with a Leica SP8 confocal system using 63X/1.40 and 2.5 zoom oil. Z-stacks of 0.4-micron interval were captured. The cell apicobasal height was calculated manually by analysing the orthogonal sections in Fiji ImageJ.

### Cell Apical area measurement

Discs expressing DE-cadherin GFP or stained with E-cadherin were used to measure the cell apical area. The wing and haltere discs were mounted with an apical side facing the objective. Apical sections of DP cells were acquired with Leica SP8 63x/1.40 Oil, 2.5 zoom and 0.4 micron step size. To analyse the cell apical area, at least three ROIs of 15X15 microns within the pouch region of each disc were analysed. 4 slices were Z-projected to accurately measure the cell size as the pouch has a curvature. For all images, Gaussian blur is set to 0.5, and the background is subtracted to rolling 50, to get clear cell boundaries for segmentation. The cells were then segmented using an ImageJ plugin - Tissue analyzer (Aigouy et al., 2016). Corresponding pixels were converted into microns later.

### Measurement of fluorescent intensities

To quantify the apical fluorescent intensities of Actin and Myosin-II in the DP cells of the imaginal wing and haltere discs, at least three ROIs of 5X5 micron square were randomly selected within the pouch region. the first 12 apical confocal z-sections of DP acquired with 0.4-micron interval were carefully selected and projected using the sum of slices function in ImageJ. Within the 5x5 ROIs, smaller ROIs of 1-micron squares were placed on cell membranes randomly, and the mean intensity was measured. Fluorescence intensity at these edges was calculated manually using Fiji with an in-house macro. To determine the average fluorescence for each disc, mean intensity values along each disc were normalised relative to the average total intensity fluorescence observed in control haltere discs.

### Cell volume Analysis

Three 15X15 um2 ROIs were chosen arbitrarily from the pouch region of confocal Z-stacks of both the wing and haltere discs that were stained for F-Actin. The mean cell height of Disc proper epithelial cells was obtained by averaging over five measurements taken for each ROI. The number of cells in each ROI was estimated using the Tissue Analyser plugin in ImageJ and corrected manually for boundary cells. The mean cell volume of the Disc proper epithelial cells for each ROI was obtained by multiplying the mean cell height of epithelial cells with the area of the ROI (i.e. 225 um^2^), to obtain the pouch volume of epithelial cells of the ROI. This product was divided with the number of cells in each ROI to obtain the mean cell volume, which was then averaged over three independent ROIs to report the volume of the disc cells.

### Scanning electron microscopy

Scanning electron microscopy (SEM) was carried out on Carl Zeiss EVO LS10 Scanning Electron Microscope Zeiss using Axiovision 4.8.2 software to operate the microscope and for image analysis. Fresh samples of flies were cleaned with 70% ethanol, flash-freezed and were directly used for imaging.

### Adult cuticle preparation/Quantification of the haltere capitellum size

Female adult flies of specified genotypes were collected and sequentially dehydrated in 10%, 50%, 70%, and 100% ethanol for 1 hour each. The flies were then left in clove oil overnight. The wing and halteres were dissected and mounted in DPX mountant (for the quantification of the haltere capitellum size).

### Measurement of trichome density

Bright-field Images of adult haltere cuticles were taken using a Zeiss Apotome microscope at 40X magnification. Ten wing blade or haltere capitellum per each genotype were analysed. The number of trichomes in three different ROIs of 30X30 micron square area per sample was manually estimated using Image J software.

### Mathematical Modelling of developmental transitions during wing and haltere morphogenesis

The lateral section vertex model is similar to the standard apical 2D vertex model. Each cell is formed of four vertices that are connected with four edges (Fig 6A). There are three types of cell boundaries, apical, lateral and basal, with contractilities or line tensions given by *λ*_*a*_, *λ*_*b*_, *λ*_*l*_. In our model, the apical and basal boundaries also have linear springs of stiffness *k*_*a*_and *k*_*b*_, respectively, that idealize the mechanical resistance of ECM at these locations — the rest length of these springs *l*_0*a*_and *l*_0*b*_are equal to the corresponding edge lengths at the start of the simulation. The preferred size of individual cell is *A*_0*c*_ and the corresponding stiffness is *K*_cell_. In the case of the haltere, there is a central lumen that is idealised as a single cell of stiffness *K*_lum_and preferred area *A*_0*l*_ (Fig 6A, central blue cell). The work function of the system is given by a simple modification of the standard vertex model (Alt et al., 2017; Farhadifar et al., 2007)

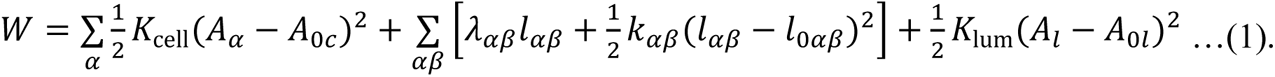

Here, the index *α* goes over all over the cells and the index *αβ* correspond to an edge that is shared between cells *α* and *β* (the exterior is labelled at zero). Note that, as discussed above, lumen is also considered to be a cell. The dynamics of lumen can be controlled by increasing or decreasing the preferred lumen size *A*_0*l*_(*t*) as a function of time. For the case of wing, a thin layer of cells in the model (Fig 6B top, thin blue layer of cells) is interpreted to be the cohesive layer between the top and bottom layers of the wing epithelium. In this case, a contribution Δ*W* from these cells is added to the work function *W* above as

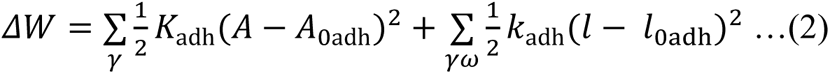

where *K*_adh_ and *A*_0adh_are the area modulus and preferred area of the cohesive cells, while *k*_adh_and *l*_0adh_are the stiffness and rest length of the springs along the vertical edges that resist separation between the top and the bottom layers of the wing (Fig 6B). The summation goes over the cells γ and the shared edges γω in this adhesive layer. This is a simplistic but a reasonably effective way of implementing the adhesion between the top and the bottom layer.

We start with initial geometries as shown in Fig 6B-D (top) such that the initial values 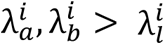 making columnar geometry as the preferred cell shape (Hannezo et al., 2014), while the rest of forcing terms are as given in Eq. 1 (and Eq. 2 for wing, Fig 6B). The system is in mechanical equilibrium in this configuration. Starting from this configuration in which the system is in mechanical equilibrium, the line tensions λ(*t*) are gradually modified with time *t* as

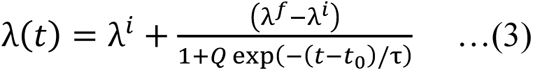

such that the final values become 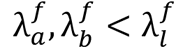 and the individual cells prefer to have a cuboidal geometry. Here *t*_0_, τ are the time scales implemented to ensure gradual modifications of line tensions. Simultaneously, the basal and apical spring stiffnesses are also gradually degraded to zero as

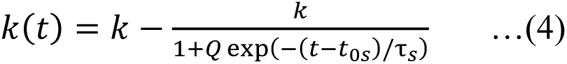

As in the case of line tensions, *t*_0*s*_, τ_*s*_ are the time-scales for gradual ECM degradation. In the case of haltere-like geometries (Fig 6C-E), the preferred lumen size *A*_0*l*_ (blue region) is made to expand from its initial value with a constant rate *dA*_0*l*_/*dt* = *r*. Finally, the position ***r***_*v*_ of any vertex *v* evolves as per the standard vertex model dynamics as η*d****r***_*v*_/*dt* = − ∂*W*/∂***r***_*v*_ (Δ*W*is also used in the case of wing), where the right-hand and the left-hand sides, respectively, provide the effective external friction force and total internal forces acting on the vertex. The equations are numerically integrated using Euler’s method with time step Δ*t* for a total duration of *T*_dur_. The values of parameters used for the simulations are presented in Table 1.

### Statistical analysis

Statistical analysis of data was performed using the GraphPad Prism version 8.4.3 software. The average and standard deviation were represented in the graphs. Unpaired t-tests were used to determine whether distribution means are significantly different between wing and haltere experiments in normally distributed data. One way ANOVA was used to determine the significance when more than two samples were analysed. Mann-Whitney test and Kruskal-Wallis test were used in case of not normally distributed data. All the tests were done using GraphPad Prism 8.4.3 software. ns, not significant, *p < 0.05, **p < 0.01, ***p < 0.001, ****p < 0.0001.

## References

Agrawal, P., Habib, F., Yelagandula, R., & Shashidhara, L. S. (2011). Genome-level identification of targets of Hox protein Ultrabithorax in Drosophila: novel mechanisms for target selection. Scientific Reports, 1(1), 205. 10.1038/srep00205

Aigouy, B., Umetsu, D., & Eaton, S. (2016). Segmentation and Quantitative Analysis of Epithelial Tissues. Methods in Molecular Biology (Clifton, N.J.), 1478, 227–239. 10.1007/978-1-4939-6371-3_13

Aldaz, S., Escudero, L. M., & Freeman, M. (2010). Live imaging of *Drosophila* imaginal disc development. Proceedings of the National Academy of Sciences, 107(32), 14217–14222. 10.1073/pnas.1008623107

Alt, S., Ganguly, P., & Salbreux, G. (2017). Vertex models: from cell mechanics to tissue morphogenesis. Philosophical Transactions of the Royal Society B: Biological Sciences, 372(1720), 20150520. 10.1098/rstb.2015.0520

Bardet, P.-L., Guirao, B., Paoletti, C., Serman, F., Léopold, V., Bosveld, F., Goya, Y., Mirouse, V., Graner, F., & Bellaïche, Y. (2013). PTEN Controls Junction Lengthening and Stability during Cell Rearrangement in Epithelial Tissue. Developmental Cell, 25(5), 534–546. 10.1016/j.devcel.2013.04.020

Cabrera, C. V., Botas, J., & Garcia-Bellido, A. (1985). Distribution of Ultrabithorax proteins in mutants of Drosophila bithorax complex and its transregulatory genes. Nature, 318(6046), 569–571. 10.1038/318569a0

Castelli-Gair, J. E., Micol, J. L., & García-Bellido, A. (1990). Transvection in the Drosophila Ultrabithorax gene: a Cbx1 mutant allele induces ectopic expression of a normal allele in trans. Genetics, 126(1), 177–184. 10.1093/genetics/126.1.177

Charroux, B., Freeman, M., Kerridge, S., & Baonza, A. (2006). Atrophin contributes to the negative regulation of epidermal growth factor receptor signaling in Drosophila. Developmental Biology, 291(2), 278–290. 10.1016/j.ydbio.2005.12.012

Chen, C.-Y., Chen, J., He, L., & Stiles, B. L. (2018). PTEN: Tumor Suppressor and Metabolic Regulator. Frontiers in Endocrinology, 9. 10.3389/fendo.2018.00338

Crickmore, M. A., & Mann, R. S. (2006). Hox Control of Organ Size by Regulation of Morphogen Production and Mobility. Science, 313(5783), 63–68. 10.1126/science.1128650

Dawes-Hoang, R. E., Parmar, K. M., Christiansen, A. E., Phelps, C. B., Brand, A. H., & Wieschaus, E. F. (2005). *folded gastrulation*, cell shape change and the control of myosin localization. Development, 132(18), 4165–4178. 10.1242/dev.01938

De las Heras, J. M., García-Cortés, C., Foronda, D., Pastor-Pareja, J. C., Shashidhara, L. S., & Sánchez-Herrero, E. (2018). The Drosophila Hox gene Ultrabithorax controls appendage shape by regulating extracellular matrix dynamics. Development, 145(13). 10.1242/dev.161844

de Navas, L., Foronda, D., Suzanne, M., & Sánchez-Herrero, E. (2006). A simple and efficient method to identify replacements of P-lacZ by P-Gal4 lines allows obtaining Gal4 insertions in the bithorax complex of Drosophila. Mechanisms of Development, 123(11), 860–867. 10.1016/j.mod.2006.07.010

Diaz-de-la-Loza, M.-C., Loker, R., Mann, R. S., & Thompson, B. J. (2020). Control of tissue morphogenesis by the HOX gene *Ultrabithorax*. Development, 147(5). 10.1242/dev.184564

Diaz-de-la-Loza, M.-C., Ray, R. P., Ganguly, P. S., Alt, S., Davis, J. R., Hoppe, A., Tapon, N., Salbreux, G., & Thompson, B. J. (2018). Apical and Basal Matrix Remodeling Control Epithelial Morphogenesis. Developmental Cell, 46(1), 23–39.e5. 10.1016/j.devcel.2018.06.006

Díaz-de-la-Loza, M.-C., & Stramer, B. M. (2024). The extracellular matrix in tissue morphogenesis: No longer a backseat driver. Cells & Development, 177, 203883. 10.1016/j.cdev.2023.203883

Domínguez-Giménez, P., Brown, N. H., & Martín-Bermudo, M. D. (2007). Integrin-ECM interactions regulate the changes in cell shape driving the morphogenesis of the *Drosophila* wing epithelium. Journal of Cell Science, 120(6), 1061–1071. 10.1242/jcs.03404

Erkner, A., Roure, A., Charroux, B., Delaage, M., Holway, N., Coré, N., Vola, C., Angelats, C., Pagès, F., Fasano, L., & Kerridge, S. (2002). Grunge, related to human Atrophin-like proteins, has multiple functions in *Drosophila* development. Development, 129(5), 1119–1129. 10.1242/dev.129.5.1119

Farhadifar, R., Röper, J.-C., Aigouy, B., Eaton, S., & Jülicher, F. (2007). The Influence of Cell Mechanics, Cell-Cell Interactions, and Proliferation on Epithelial Packing. Current Biology, 17(24), 2095–2104. 10.1016/j.cub.2007.11.049

Fernandez-Gonzalez, R., Simoes, S. de M., Röper, J.-C., Eaton, S., & Zallen, J. A. (2009). Myosin II Dynamics Are Regulated by Tension in Intercalating Cells. Developmental Cell, 17(5), 736–743. 10.1016/j.devcel.2009.09.003

Gao, X., Neufeld, T. P., & Pan, D. (2000). Drosophila PTEN Regulates Cell Growth and Proliferation through PI3K-Dependent and -Independent Pathways. Dev Biol, 221(2), 404–418. 10.1006/dbio.2000.9680

Giraud, G., Paul, R., Duffraisse, M., Khan, S., Shashidhara, L. S., & Merabet, S. (2021). Developmental Robustness: The Haltere Case in Drosophila. Frontiers in Cell and Developmental Biology, 9. 10.3389/fcell.2021.713282

Goberdhan, D. C. I., Paricio, N., Goodman, E. C., Mlodzik, M., & Wilson, C. (1999). Drosophila tumor suppressor PTEN controls cell size and number by antagonizing the Chico/PI3-kinase signaling pathway. Genes & Development, 13(24), 3244–3258. 10.1101/gad.13.24.3244

Gokhale, R. H., & Shingleton, A. W. (2015). Size control: the developmental physiology of body and organ size regulation. WIREs Developmental Biology, 4(4), 335–356. 10.1002/wdev.181

Groth, C., Vaid, P., Khatpe, A., Prashali, N., Ahiya, A., Andrejeva, D., Chakladar, M., Nagarkar, S., Paul, R., Kelkar, D., Eichenlaub, T., Herranz, H., Sridhar, T. S., Cohen, S. M., & Shashidhara, L. S. (2020). Genome-Wide Screen for Context-Dependent Tumor Suppressors Identified Using in Vivo Models for Neoplasia in Drosophila. G3 (Bethesda), 10(9), 2999–3008. 10.1534/g3.120.401545

Hannezo, E., Prost, J., & Joanny, J.-F. (2014). Theory of epithelial sheet morphology in three dimensions. Proceedings of the National Academy of Sciences, 111(1), 27–32. 10.1073/pnas.1312076111

Hansen, C. G., Moroishi, T., & Guan, K.-L. (2015). YAP and TAZ: a nexus for Hippo signaling and beyond. Trends in Cell Biology, 25(9), 499–513. 10.1016/j.tcb.2015.05.002

Harmansa, S., Erlich, A., Eloy, C., Zurlo, G., & Lecuit, T. (2023). Growth anisotropy of the extracellular matrix shapes a developing organ. Nature Communications, 14(1), 1220. 10.1038/s41467-023-36739-y

Heer, N. C., & Martin, A. C. (2017). Tension, contraction and tissue morphogenesis. Development, 144(23), 4249–4260. 10.1242/dev.151282

Heisenberg, C.-P., & Bellaïche, Y. (2013). Forces in Tissue Morphogenesis and Patterning. Cell, 153(5), 948–962. 10.1016/j.cell.2013.05.008

Huang, J., Wu, S., Barrera, J., Matthews, K., & Pan, D. (2005). The Hippo Signaling Pathway Coordinately Regulates Cell Proliferation and Apoptosis by Inactivating Yorkie, the Drosophila Homolog of YAP. Cell, 122(3), 421–434. 10.1016/j.cell.2005.06.007

Irvine, K. D., & Harvey, K. F. (2015). Control of Organ Growth by Patterning and Hippo Signaling in *Drosophila*. Cold Spring Harbor Perspectives in Biology, 7(6), a019224. 10.1101/cshperspect.a019224

Katsumi, A., Orr, A. W., Tzima, E., & Schwartz, M. A. (2004). Integrins in Mechanotransduction. Journal of Biological Chemistry, 279(13), 12001–12004. 10.1074/jbc.R300038200

Khan, S., Dilsha, C., & Shashidhara, L. S. (2020). Haltere development in D. melanogaster: implications for the evolution of appendage size, shape and function. Int J Dev Biol, 64(1-2–3), 159–165. 10.1387/ijdb.190133LS

Kondo, T., & Hayashi, S. (2015). Mechanisms of cell height changes that mediate epithelial invagination. Development, Growth & Differentiation, 57(4), 313–323. 10.1111/dgd.12224

Kozyrina, A. N., Piskova, T., & Di Russo, J. (2020). Mechanobiology of Epithelia From the Perspective of Extracellular Matrix Heterogeneity. Frontiers in Bioengineering and Biotechnology, 8. 10.3389/fbioe.2020.596599

Lewis, E. B. (1978). A Gene Complex Controlling Segmentation in Drosophila. In Genes, Development, and Cancer (pp. 229–242). Springer Netherlands. 10.1007/978-1-4020-6345-9_10

Liu, P., Guo, Y., Xu, W., Song, S., Li, X., Wang, X., Lu, J., Guo, X., Richardson, H. E., & Ma, X. (2022). Ptp61F integrates Hippo, TOR, and actomyosin pathways to control three-dimensional organ size. Cell Reports, 41(7), 111640. 10.1016/j.celrep.2022.111640

Luciano, M., Versaevel, M., Vercruysse, E., Procès, A., Kalukula, Y., Remson, A., Deridoux, A., & Gabriele, S. (2022). Appreciating the role of cell shape changes in the mechanobiology of epithelial tissues. Biophysics Reviews, 3(1). 10.1063/5.0074317

Makhijani, K., Kalyani, C., Srividya, T., & Shashidhara, L. S. (2007). Modulation of Decapentaplegic gradient during haltere specification in Drosophila. Dev Biol, 302(1), 243–255. 10.1016/j.ydbio.2006.09.029

Martin, A. C., & Goldstein, B. (2014). Apical constriction: themes and variations on a cellular mechanism driving morphogenesis. Development, 141(10), 1987–1998. 10.1242/dev.102228

Martin, P. F. (1982). Direct determination of the growth rate of Drosophila imaginal discs. Journal of Experimental Zoology, 222(1), 97–102. 10.1002/jez.1402220113

Miao, H., & Blankenship, J. T. (2020). The pulse of morphogenesis: actomyosin dynamics and regulation in epithelia. Development, 147(17). 10.1242/dev.186502

Misra, J. R., & Irvine, K. D. (2018). The Hippo Signaling Network and Its Biological Functions. Annual Review of Genetics, 52(1), 65–87. 10.1146/annurev-genet-120417-031621

Mohit, P., Makhijani, K., Madhavi, M. B., Bharathi, V., Lal, A., Sirdesai, G., Reddy, V. R., Ramesh, P., Kannan, R., Dhawan, J., & Shashidhara, L. S. (2006). Modulation of AP and DV signaling pathways by the homeotic gene Ultrabithorax during haltere development in Drosophila. Developmental Biology, 291(2), 356–367. 10.1016/j.ydbio.2005.12.022

Morata G, Garcia-Bellido A. Developmental analysis of some mutants of the bithorax system of Drosophila. Wilehm Roux Arch Dev Biol. 1976 Jun;179(2):125–143. doi: 10.1007/BF00848298.

Nagarkar, S., Wasnik, R., Govada, P., Cohen, S., & Shashidhara, L. S. (2020). Promoter Proximal Pausing Limits Tumorous Growth Induced by the Yki Transcription Factor in *Drosophila*. Genetics, 216(1), 67–77. 10.1534/genetics.120.303419

Nelson, C. M., & Gleghorn, J. P. (2012). Sculpting Organs: Mechanical Regulation of Tissue Development. Annual Review of Biomedical Engineering, 14(1), 129–154. 10.1146/annurev-bioeng-071811-150043

Pallavi, S. K., Kannan, R., & Shashidhara, L. S. (2006). Negative regulation of Egfr/Ras pathway by Ultrabithorax during haltere development in Drosophila. Dev Biol, 296(2), 340–352. 10.1016/j.ydbio.2006.05.035

Pallavi, S. K., & Shashidhara, L. S. (2003). Egfr/Ras pathway mediates interactions between peripodial and disc proper cells in Drosophila wing discs. Development, 130(20), 4931–4941. 10.1242/dev.00719

Pan, D. (2010). The Hippo Signaling Pathway in Development and Cancer. Developmental Cell, 19(4), 491–505. 10.1016/j.devcel.2010.09.011

Pavlopoulos, A., & Akam, M. (2011). Hox gene *Ultrabithorax* regulates distinct sets of target genes at successive stages of *Drosophila* haltere morphogenesis. Proceedings of the National Academy of Sciences, 108(7), 2855–2860. 10.1073/pnas.1015077108

Polyakov, O., He, B., Swan, M., Shaevitz, J. W., Kaschube, M., & Wieschaus, E. (2014). Passive Mechanical Forces Control Cell-Shape Change during Drosophila Ventral Furrow Formation. Biophysical Journal, 107(4), 998–1010. 10.1016/j.bpj.2014.07.013

Prasad, M., Bajpai, R., & Shashidhara, L. S. (2003). Regulation of Wingless and Vestigial expression in wing and haltere discs of *Drosophila*. Development, 130(8), 1537–1547. 10.1242/dev.00393

Rauzi, M., Lenne, P.-F., & Lecuit, T. (2010). Planar polarized actomyosin contractile flows control epithelial junction remodelling. Nature, 468(7327), 1110–1114. 10.1038/nature09566

Roch, F., & Akam, M. (2000). Ultrabithorax and the control of cell morphology in Drosophila halteres. Development, 127(1), 97–107. 10.1242/dev.127.1.97

Schwartz, M. A. (2010). Integrins and Extracellular Matrix in Mechanotransduction. Cold Spring Harbor Perspectives in Biology, 2(12), a005066–a005066. 10.1101/cshperspect.a005066

Shashidhara, L. S., Agrawal, N., Bajpai, R., Bharathi, V., & Sinha, P. (1999). Negative Regulation of Dorsoventral Signaling by the Homeotic Gene Ultrabithorax during Haltere Development in Drosophila. Developmental Biology, 212(2), 491–502. 10.1006/dbio.1999.9341

Singh, S., Sanchez-Herrero, E., & Shashidhara, L. S. (2015). Critical role for Fat/Hippo and IIS/Akt pathways downstream of Ultrabithorax during haltere specification in Drosophila. Mech Dev, 138 *Pt* *2*, 198–209. 10.1016/j.mod.2015.07.017

Song, S., Herranz, H., & Cohen, S. M. (2017). The chromatin remodeling BAP complex limits tumor promoting activity of the Hippo pathway effector Yki to prevent neoplastic transformation in *Drosophila* epithelia. Disease Models & Mechanisms. 10.1242/dmm.030122

Sui, L., Alt, S., Weigert, M., Dye, N., Eaton, S., Jug, F., Myers, E. W., Jülicher, F., Salbreux, G., & Dahmann, C. (2018). Differential lateral and basal tension drive folding of Drosophila wing discs through two distinct mechanisms. Nature Communications, 9(1), 4620. 10.1038/s41467-018-06497-3

Sun, T., Song, Y., Teng, D., Chen, Y., Dai, J., Ma, M., Zhang, W., & Pastor-Pareja, J. C. (2021). Atypical laminin spots and pull-generated microtubule-actin projections mediate Drosophila wing adhesion. Cell Reports, 36(10), 109667. 10.1016/j.celrep.2021.109667

Sun, Z., Guo, S. S., & Fässler, R. (2016). Integrin-mediated mechanotransduction. Journal of Cell Biology, 215(4), 445–456. 10.1083/jcb.201609037

Tozluoǧlu, M., & Mao, Y. (2020). On folding morphogenesis, a mechanical problem. Philosophical Transactions of the Royal Society B: Biological Sciences, 375(1809), 20190564. 10.1098/rstb.2019.0564

Vasquez, C. G., Tworoger, M., & Martin, A. C. (2014). Dynamic myosin phosphorylation regulates contractile pulses and tissue integrity during epithelial morphogenesis. Journal of Cell Biology, 206(3), 435–450. 10.1083/jcb.201402004

Wang, L., Charroux, B., Kerridge, S., & Tsai, C. (2008). Atrophin recruits HDAC1/2 and G9a to modify histone H3K9 and to determine cell fates. EMBO Reports, 9(6), 555–562. 10.1038/embor.2008.67

Wang, L., Rajan, H., Pitman, J. L., McKeown, M., & Tsai, C.-C. (2006). Histone deacetylase-associating Atrophin proteins are nuclear receptor corepressors. Genes & Development, 20(5), 525–530. 10.1101/gad.1393506

Wang, L., & Tsai, C.-C. (2008). Atrophin Proteins: An Overview of a New Class of Nuclear Receptor Corepressors. Nuclear Receptor Signaling, 6(1), nrs.06009. 10.1621/nrs.06009

Weatherbee, S. D., Halder, G., Kim, J., Hudson, A., & Carroll, S. (1998). Ultrabithorax regulates genes at several levels of the wing-patterning hierarchy to shape the development of the *Drosophila* haltere. Genes & Development, 12(10), 1474–1482. 10.1101/gad.12.10.1474

White, R. A. H., & Akam, M. E. (1985). Contrabithorax mutations cause inappropriate expression of Ultrabithorax products in Drosophila. Nature, 318(6046), 567–569. 10.1038/318567a0

White, R. A. H., & Wilcox, M. (1985). Regulation of the distribution of Ultrabithorax proteins in Drosophila. Nature, 318(6046), 563–567. 10.1038/318563a0

Wittkorn, E., Sarkar, A., Garcia, K., Kango-Singh, M., & Singh, A. (2015). The Hippo pathway effector Yki downregulates Wg signaling to promote retinal differentiation in the *Drosophila* eye. Development, 142(11), 2002–2013. 10.1242/dev.117358

Yeung, K., Boija, A., Karlsson, E., Holmqvist, P.-H., Tsatskis, Y., Nisoli, I., Yap, D., Lorzadeh, A., Moksa, M., Hirst, M., Aparicio, S., Fanto, M., Stenberg, P., Mannervik, M., & McNeill, H. (2017). Atrophin controls developmental signaling pathways via interactions with Trithorax-like. ELife, 6. 10.7554/eLife.23084

Zartman, J. J., & Shvartsman, S. Y. (2010). Unit Operations of Tissue Development: Epithelial Folding. Annual Review of Chemical and Biomolecular Engineering, 1(1), 231–246. 10.1146/annurev-chembioeng-073009-100919

Zheng, Y., & Pan, D. (2019). The Hippo Signaling Pathway in Development and Disease. Developmental Cell, 50(3), 264–282. 10.1016/j.devcel.2019.06.003

