## Supplementary Figures for "Differences in Cellular mechanics and ECM dynamics shape differential development of wing and haltere in *Drosophila*"

Supplementary Figures:
S1

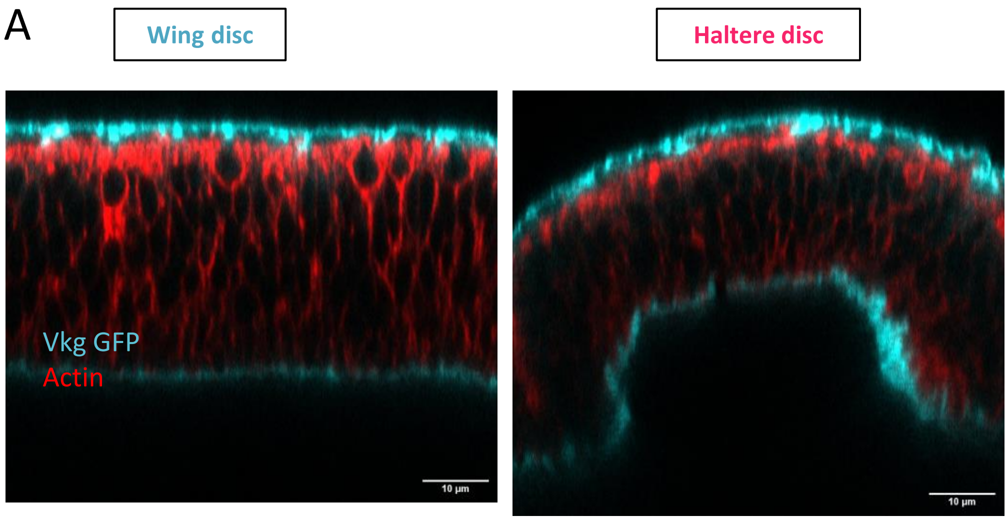


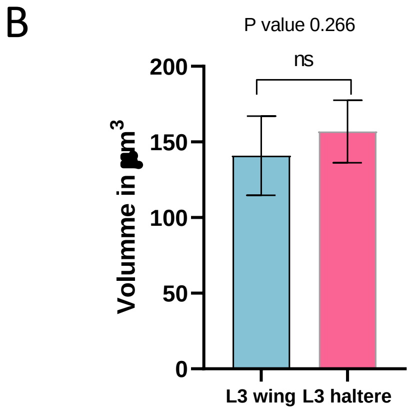

S2

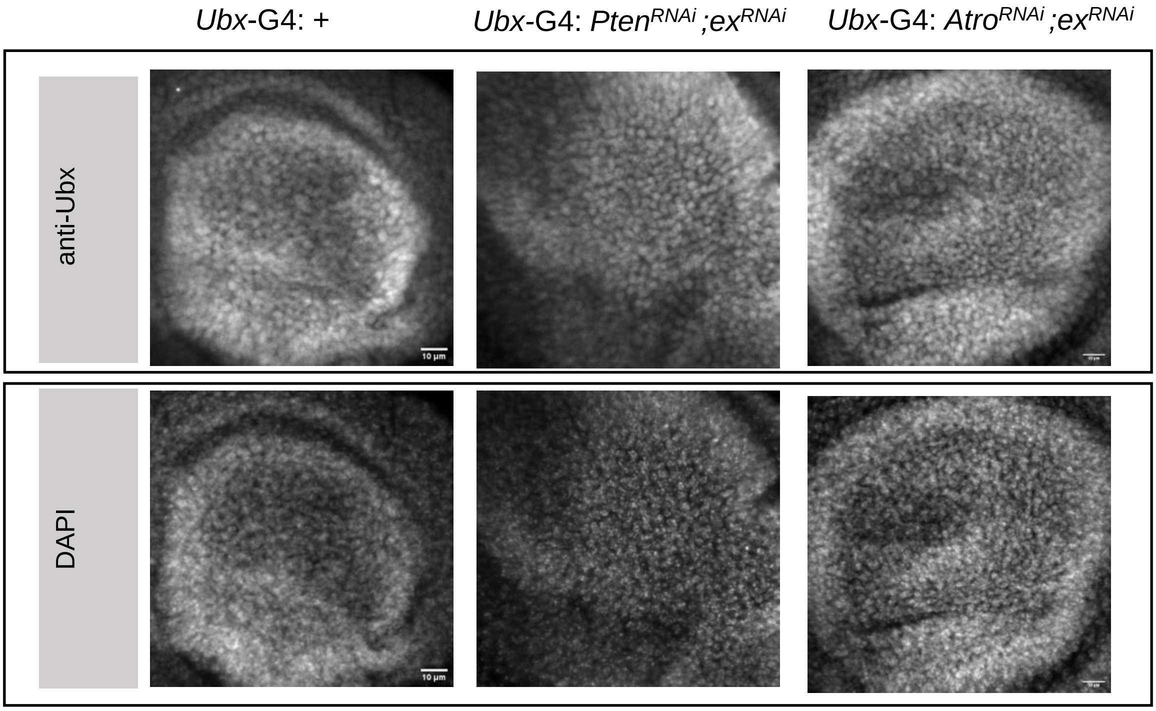
*Ubx*-G4: +
*Ubx*-G4: *Pten*$^{RNAi}$;*ex*$^{RNAi}$
*Ubx*-G4: *Atro*$^{RNAi}$;*ex*$^{RNAi}$
anti-Ubx
DAPI
10 µm

S3

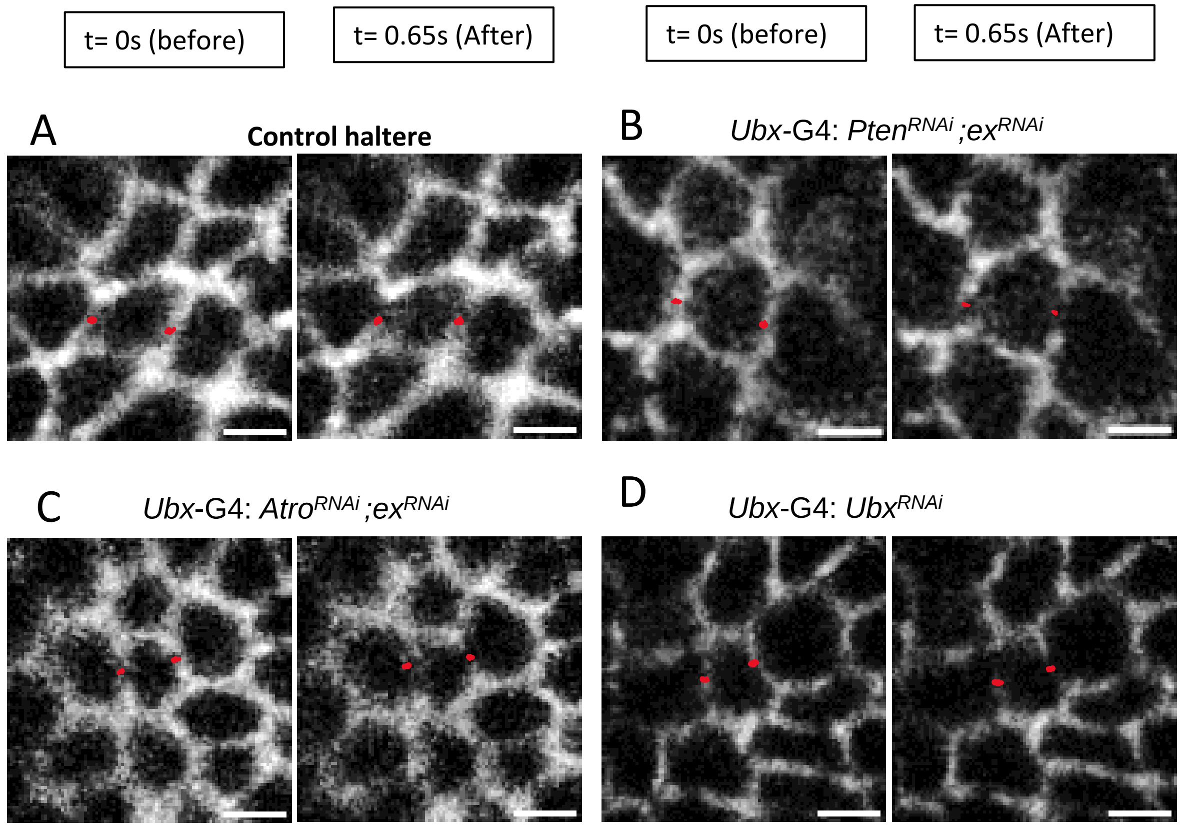
t= 0s (before)
t= 0.65s (After)
t= 0s (before)
t= 0.65s (After)
A
Control haltere
B
Ubx-G4: Pten^RNAi ;ex^RNAi
C
Ubx-G4: Atro^RNAi ;ex^RNAi
D
Ubx-G4: Ubx^RNAi

S4

A *nub* G4>UAS *Timp*

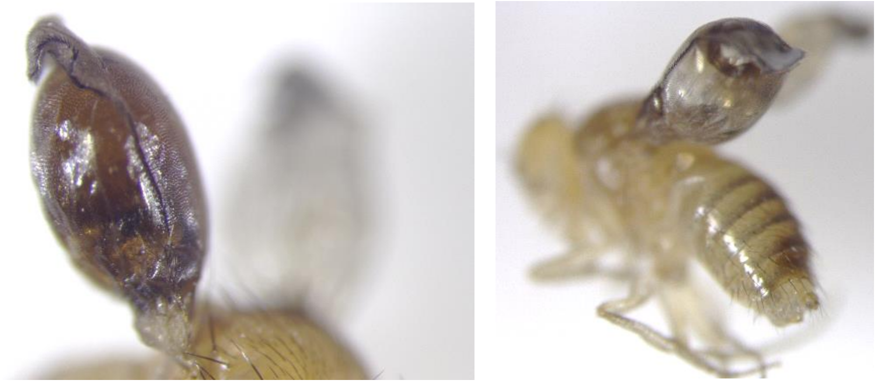

B *nub* G4>*If*$^{RNAi}$

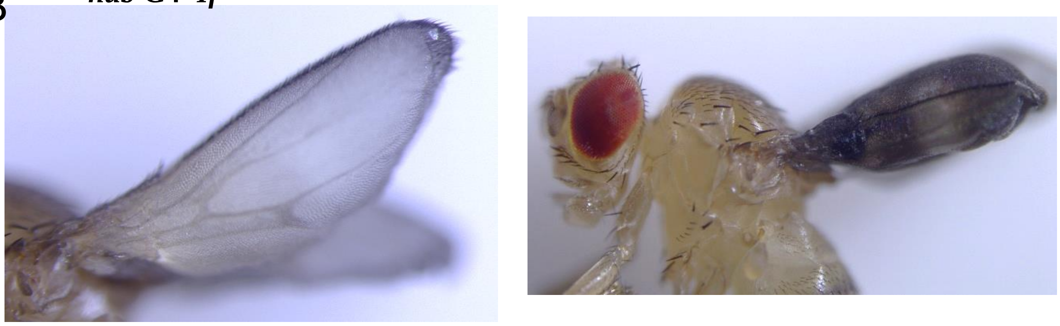

C *Ubx*-G4: *Atro*$^{RNAi}$ ;*ex*$^{RNAi}$

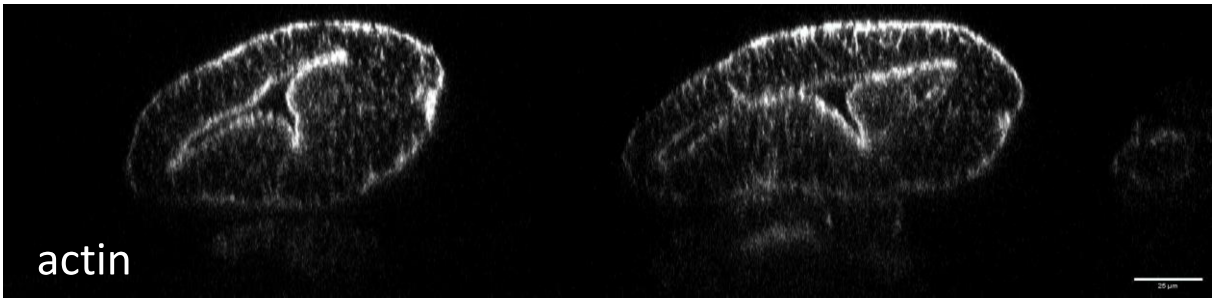
