## Supplementary Information for "Differences in Cellular mechanics and ECM dynamics shape differential development of wing and haltere in *Drosophila*"

#### Supplementary Figures

**S1**: A.Wing and haltere discs stained for F-actin and GFP to improve Vkg GFP signal. Actin signal spans throughout the length of both wing and haltere disc columnar cells. Scale bar, 10 microns.
B. Bar Plot showing the volume approximation between L3 wing and haltere discs. The cellular volume of wing and haltere discs at L3 were comparable. n=6 discs, Student's t-test.

**S2**: Haltere discs stained for Ubx in *Pten*$^{RNAi}$*; ex*$^{RNAi}$ and *Atro*$^{RNAi}$*; ex*$^{RNAi}$ mutants. There is no significant reduction in Ubx expression levels were observed within these haltere discs. DAPI staining was used as an internal control.

**S3.** Increased bond tension at apical cell junctions in *Atro*$^{RNAi}$*; ex*$^{RNAi}$, *Pten*$^{RNAi}$*; ex*$^{RNAi}$ and *Ubx*$^{RNAi}$ haltere discs. Haltere disc cells expressing endo E cadherin GFP before (t=0s) and after (t= 0.65s) the laser cut is shown. The red spots indicate the site of ablation. Scale bar, 2 μm.

**S4** A. Flies expressing Timp and B, *If*$^{RNAi}$, respectively in the wings show wing blistering. The wings appeared inflated and lacked DV apposition.

A. Cross-sections of *Atro*$^{RNAi}$*; ex*$^{RNAi}$ haltere discs (4-6h APF) showing 3D deformations.

#### Supplementary Videos

**Movie 1-2**: Live imaging of DE-Cadherin GFP expressing control wing (movie 1) and haltere cells (movie 2) subjected to laser ablations at apical junctions at L3. The movie is centred on the ablated cell. Wing disc cells shows higher recoil when compared to haltere cells. Scale bar, 2 μm.

**Movie 3-5**: Live imaging of DE-Cadherin GFP expressing *Pten*$^{RNAi}$*; ex*$^{RNAi}$ (movie 3), *Atro*$^{RNAi}$*; ex*$^{RNAi}$ (movie 4) and *Ubx*$^{RNAi}$ halteres (movie 5) subjected to laser ablations at apical junctions at L3 showed an increased recoil compared to the control haltere (movie 2). A The movie is centred on the ablated cell. Scale bar, 2 μm.

### Supplementary Information

**Table 1:** Values of simulation parameters

| Quantity | Value (non-dimensional) |
|---|---|
| $K_{\text{cell}}$ | 0.3 |
| $A_{0c}$ | $\approx 500 - 600$ (cell area at the start of simulation) |
| $\lambda_a, \lambda_b, \lambda_l$ | Columnar: (100,100,30), Cuboidal: (60,60,80) |
| $k_a, k_b$ | Fig 6B: (50,50), Fig 6C-6E: (100, 100) |
| $K_{\text{lum}}$ | 0.5 |
| $A_{0l}$ | blue cell area at the start of simulation |
| $K_{\text{adh}}$ | 0.5 |
| $A_{0\text{adh}}$ | blue cell area at the start of simulation (Fig. 6B) |
| $k_{\text{adh}}$ | 100 |
| $l_{0\text{adh}}$ | edge length at start of simulation (Fig. 6B) |
| $Q$ | 100 |
| $\Delta t$ | 1 |
| $\tau, \tau_0$ | Fig. 6B: (100,3000), Fig 6C-6E: (10000,6000) |
| $t_{0s}, \tau_s$ | Fig. 6B: (100,3000), Fig. 6C-6E: (15000,8000) |
| $T_{dur}$ | Fig. 6B, 6E: 39000, Fig. 6C, 6D: 120000 |
| $r$ | 0.25,0.5 |
| $\eta$ | $0.33 \times 10^{-2}$ |
